# Piperazine-based HIV-1 entry inhibitors: Massive in silico library design and screening for gp120 attachment inhibitors

**DOI:** 10.1101/330142

**Authors:** Durbis J. Castillo-Pazos, Antonio Romo-Mancillas, Joaquín Barroso-Flores

**Affiliations:** Centro Conjunto de Investigación en Química Sustentable UAEM-UNAM, Carretera Toluca - Atlacomulco Km. 14.5, Personal de la UNAM, Unidad San Cayetano, C.P. 50200 Toluca de Lerdo, México.; Laboratorio de Diseño Asistido por Computadora y Síntesis de Fármacos, Facultad de Química, Universidad Autónoma de Querétaro, Cerro de Las Campanas, s/n, Las Campanas, 76010 Santiago de Querétaro, Qro. México.

**Keywords:** HIV-1, gp120, entry inhibitors, HTVS, docking, induced-fit docking, combinatorial library, piperazine, CADD

## Abstract

HIV-1 attachment, despite being an ideal target stage to stop infection from the beginning, remains as one of the HIV lifecycle phases with less amount of designed and commercially available inhibitors. To contribute to the urgently needed discovery of new active compounds that could become part of the current highly active antiretroviral therapy, and as an attempt to explore a massive chemical space, high-throughput virtual screening of 16.3 million combinatorially generated and piperazine-cored compounds, was accomplished. Docking calculations, molecular dynamics simulations, and QSAR analyses were carried out to assess the suitability of each ligand to bind gp120 envelope glycoprotein, thus preventing it from binding to CD4 co-receptor. Ligand 255 stands out as a promising candidate to be tested beyond computational methodologies, and the 4,5,6,7-tetrahydroindole fragment is reported as a better group to bind inside the Phe43 cavity than the substituted indoles reported in the literature.

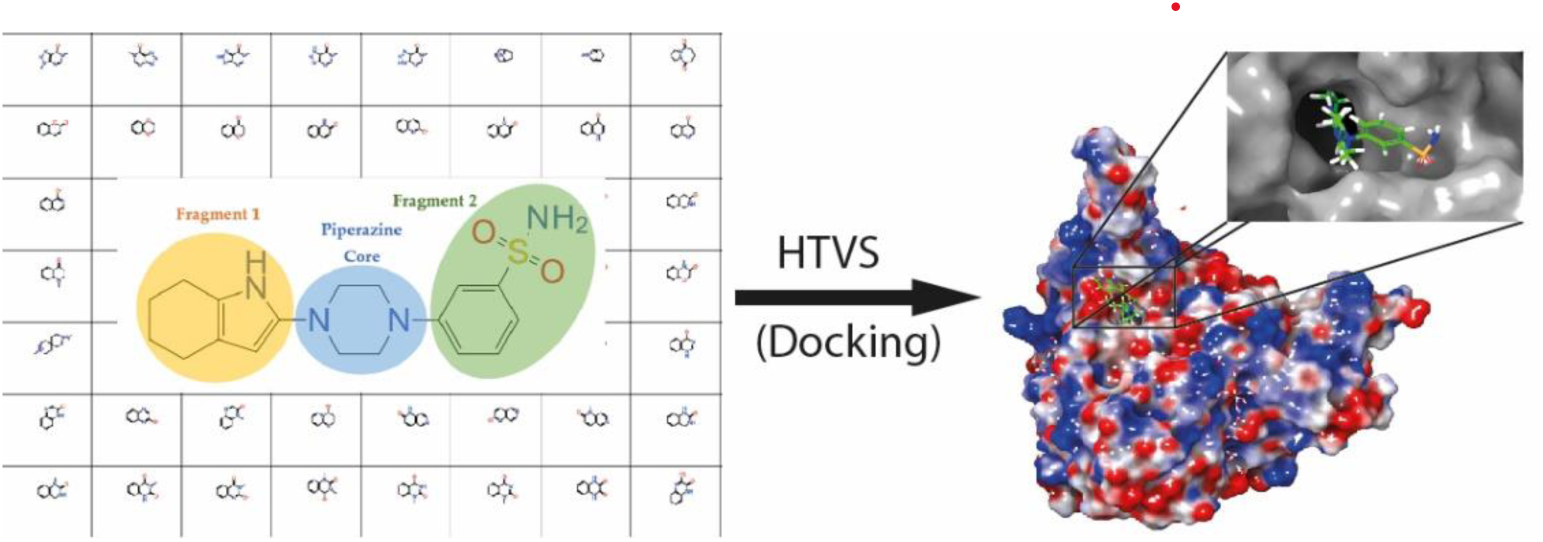

## 1. INTRODUCTION

Since its discovery in the early 1980s, the Human Immunodeficiency Virus (HIV) has attracted the attention for being the etiologic agent of Acquired Immune Deficiency Syndrome (AIDS), a global epidemic that peaked in 1999 (1) despite its probable spreading throughout the 1900s from primates to humans (2). Its high rates of recombination and mutation, persistence in the host’s body and high turnover rates allows it to elude the immune response and resist drug therapy. This high genetic diversity poses a threat to the quick effective development of treatment methods against the virus since T lymphocytes or CD4 cells are the most affected by HIV infection. These cells coordinate the action of other cells to fight infection, leading to a debilitated immune system when their count drops significantly (1).

Even though HIV-1 life cycle presents plenty of steps, enzymatic activities, and mechanisms that could be interfered by the action of drugs, very few from them have been fully exploited (3); such is the case of entry inhibitors for which a small amount of active compounds are available. By 2013, 24 drugs approved by the Food and Drug Administration (FDA) were available for treatment of HIV-1, distributed in six different classes(3): Nucleoside/Nucleotide reverse transcriptase inhibitors (NRTIs), Non-nucleoside reverse transcriptase inhibitors (NNRTIs), Integrase inhibitors (InSTIs), Protease inhibitors (PIs), and Entry inhibitors. Classified as entry inhibitors, attachment inhibitors represent a promising group of biologically active compounds, due to their capacity to stop infection from the very beginning, even though they have not been explored as much as those ligands targeting other viral proteins, such as reverse transcriptase or protease, for which most commercial drugs are designed. Among the pharmacological targets of this class of inhibitors, gp120 is of special interest thanks to its role: binding to CD4 co-receptors, marking the start of the whole HIV life cycle.

The inhibition against attachment between gp120 and CD4 receptor has been addressed with the small molecule family of BMS-378806, showing promising results in clinical assays, although it has not been yet approved. BMS-378806 alters the conformation of gp120 by binding to a pocket in which the CD4 receptor would normally enter (3).

The prototype of BMS-378806 (figure 1, compound **3**), compound **1**, was discovered thanks to a cell-based screening assay, and mechanistic studies showed that it compromised the interaction between gp120 and CD4. Since the C-4 position of the indole is lightly sensitive to the potency of the whole compound, SAR studies suggested new substitutions like the one shown in compound **2** with a fluorine atom (4). Following the improvement of the lead molecules, compound **3** showed inhibition of infection in several types of clinical isolates of the B subtype virus; however, the compound failed to achieve target exposure and its development was truncated during clinical trials (5)

**Figure 1:**
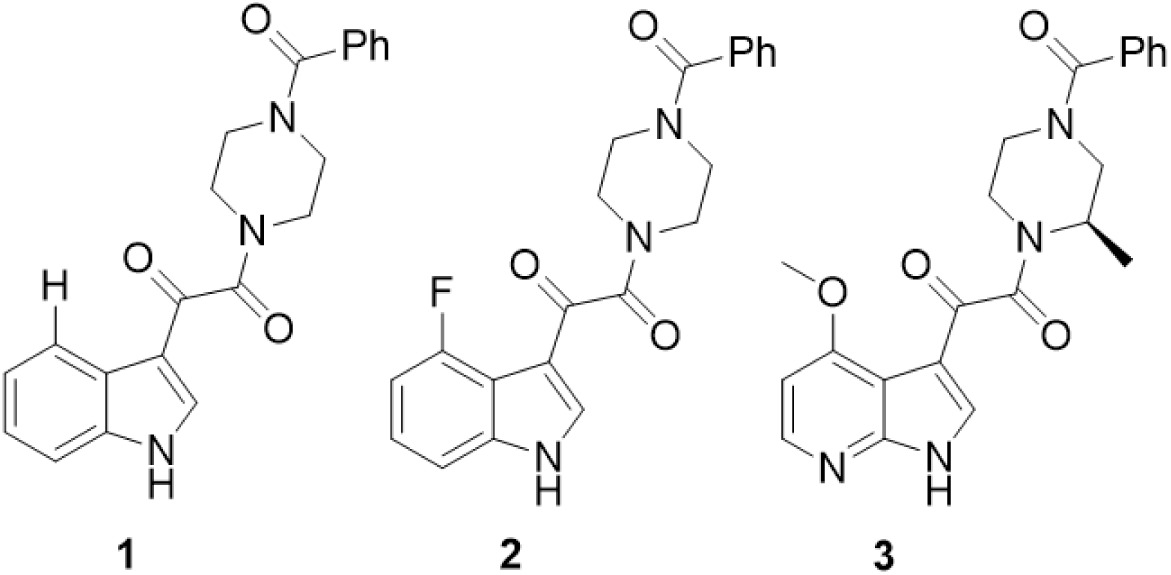
*Compounds 1 and 2, prototypes of BMS-378806 (compound 3). Compound 1 was used as the reference for Docking calculations*

The indole ring is considered the central core of these compounds because of its sensitivity to potency, offering the opportunity for structural modification. On the other hand, the piperazine ring turned out to be of critical importance for the antiviral activity (4,6), biologically and electronically. Because of its anti-HIV properties, its usual interaction at the entrance of the Phe43 cavity, and vast presence in a significant amount of anti-HIV compounds with reported activities. It is important to bear in mind that the replacement of components in existent drugs is an important strategy to afford more synthetically accessible molecules. (4).

Well established methodologies for drug discovery have been steadily replaced by mechanistic-based approaches, such as high-throughput screening, optimization of inhibitors from lead compounds, or rational drug design directly modeled on viral proteins (3). These approaches advance at a fast pace, requiring more efficient methods and techniques, and making the use of computers mandatory at all levels, from the screening of molecular libraries, to the clinical studies.

In this work, we applied several computational tools, that combined increase the possibility of finding an active compound, to the design of a molecule with high probability of showing activity against HIV gp120 envelope glycoprotein, in a 16.3 million compound library built with the software available at the time, leading to massive analysis when compared to other studies reported in the literature, none of them related to gp120-CD4 interaction (7–15)

Our main objective was to find new entry inhibitors through development and screening of a massive amount of virtually constructed molecules, capable of inhibiting the effects caused by the interaction and attachment of the Human Immunodeficiency Virus type 1, saving a considerable amount of time: while the traditional scheme of high throughput screening that is carried out every day in pharmaceutical companies takes outstanding quantities of human resources, time, and money, the *in silico* search for new active compounds performed in this work pretends to point out which ligands are worth synthesizing and testing from the beginning; thus, in this paper, we will discuss the results of our high-throughput virtual screening, the evaluation the binding ability of the top-score ligands and the prediction of the biological activity through Absorption-Distribution-Metabolism-Excretion (ADME) analysis inside the explored chemical space.

## 2. METHODS

This work is divided into three main stages: generation, screening and validation, and refinement; each one of them depending on the preceding one, with the possibility of becoming an iterative process to find molecules with higher binding affinity than those analyzed in the previous screening, included afterwards inside a consensus docking stage in the last steps of the docking funnel, using two PDB files and two different docking programs to avoid biased ranking. Below we discuss each one of the procedures included in the three main stages.

### 2.1 Protein preparation

The structure of the envelope glycoprotein gp120 was prepared with the Protein Preparation Wizard(16) included in Maestro(16). The structure was imported from the Protein Data Bank under the PDB code 1GC1(17) including substructures C (CD4 receptor), G (gp120 protein), L and H (antibodies 17B). The whole structure was preprocessed with the predetermined values at the Protein Preparation Wizard(16), filling missing chains and capping termini with the help of Prime(16), removing all water molecules beyond 5.00 Å from heteroatom groups, and generating tautomeric states of heteroatom groups at pH 7.4 ± 0.5 using Epik(16).

After substructures H and L (from antibody 17B) were removed from the workspace, along with their associated heteroatom groups and water molecules, heteroatom groups from gp120, such as *N*-acetyl-D-glucosamine (NAG), 2-(acetylamino)-2-deoxy-α-D-glucopyranose (NDG), and α-L-fucose (FUC), were left attached to the envelope protein since they are a part of its structure and do not modify the structural distribution of the Phe 43 cavity. The remaining substructures C and G (figure 2) were refined through three consecutive restrained optimizations within 0.30 Å RMSD convergence for heavy atoms at pH 7.4.

**Figure 2:**
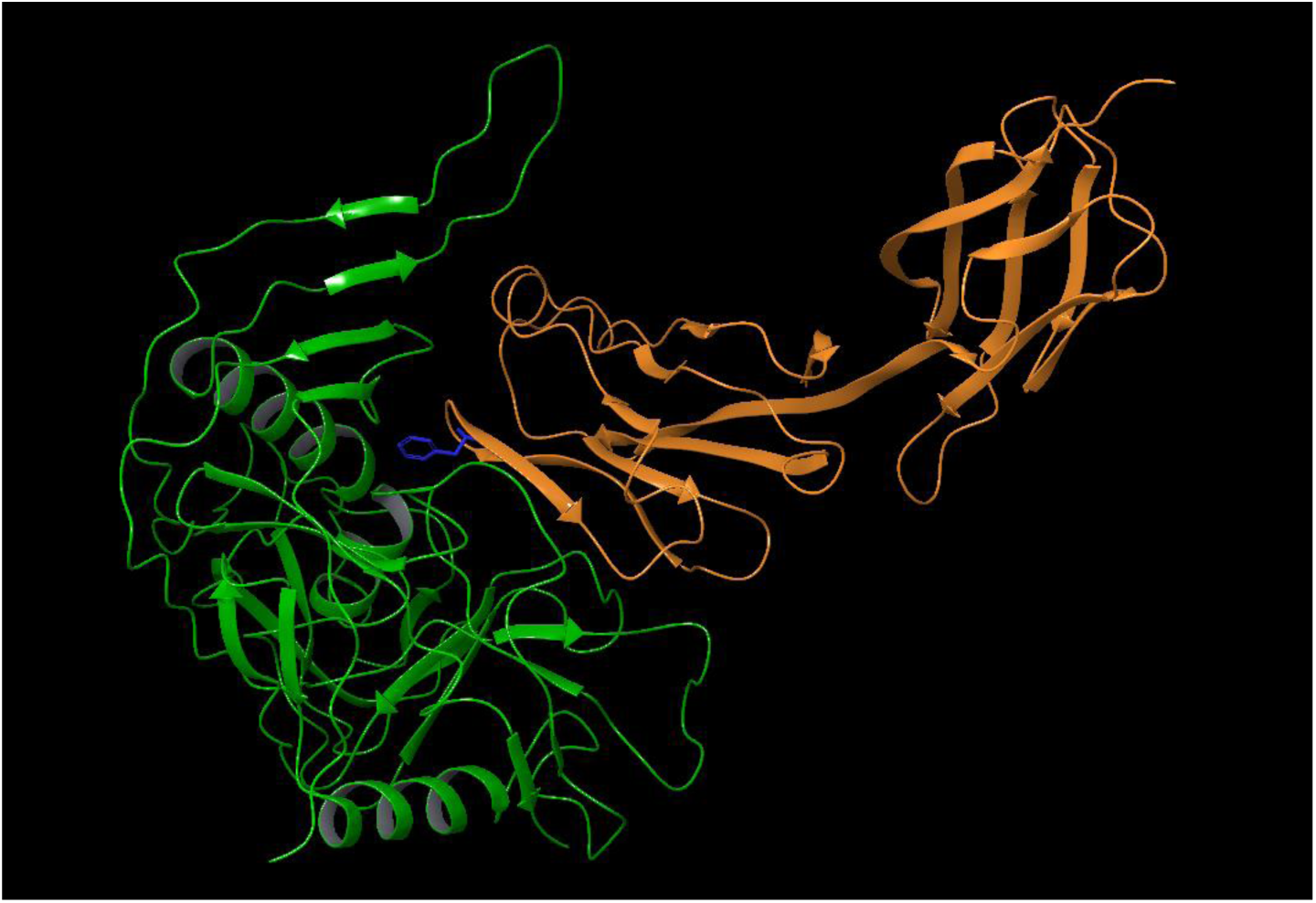
Interaction between gp120 (green) and CD4 (orange) proteins. The attachment of the virus depends on the insertion of CD4 Phe43 residue (blue) in the gp120 cavity.

**Figure 3:**
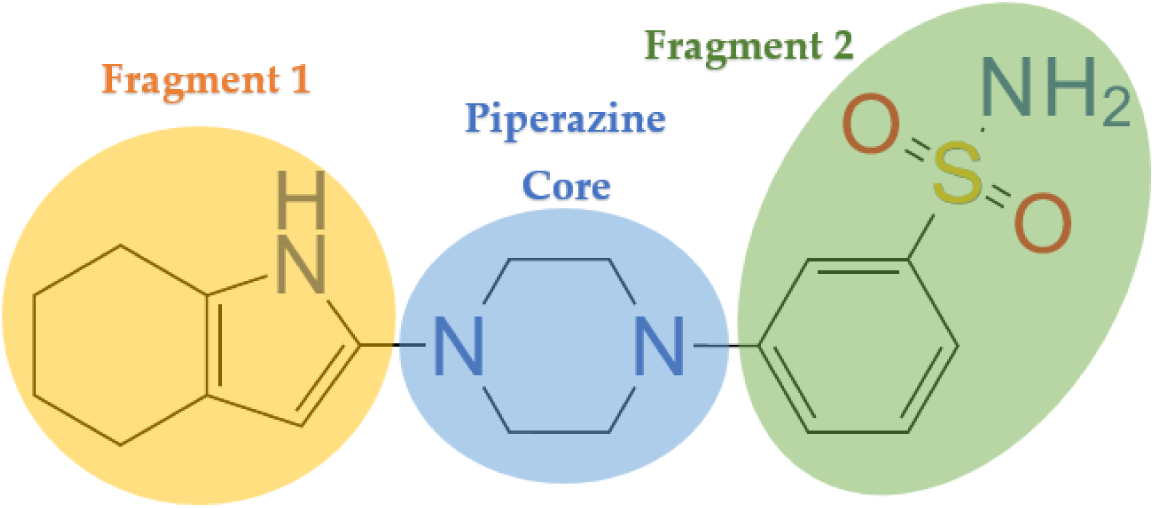
Example of a combinatorially generated molecule by Combiglide, made of three sections, a piperazine core and two different or equal molecular fragments taken out of a previously prepared database

The resulting structure was submitted to a final series of minimizations under an implicit solvent refinement process. In this case, the positions of all atoms were optimized from the substructures were minimized under a VSGB solvation model, using the automatic method provided by Prime(16), with 65 steps by iteration, and a total of 2 iterations. An RMSD gradient of 0.01 kcal/(mol Å) was considered for convergence. This process was repeated three times, until the final structure was visually checked removing the substructure C from the workspace, allowing the Phe43 cavity to be both accessible to a ligand, and suitable for the generation of a grid. Same steps were taken to prepare the gp120 structure coming from the 1G9M(18) PDB file used in the final steps of the docking stage, called consensus docking, preventing a structural bias in the docking score values.

### 2.2 Combinatorial library production and grid generation

Since piperazine was a common component among many anti-HIV compounds described in the literature, and its important role in anti-HIV activity has been identified (6), we decided to use this scaffold as the core of a series of combinatorially generated virtual compounds to perform high throughput virtual screening using Glide and its set of score functions to obtain possible hits. To generate this library, we turned to Combiglide(16) and two of its main features: Interactive Enumeration and Receptor Grid Generation.

Our fragment collection was built by importing the Glide 670-fragment database provided by Schrödinger Inc. online. A total of 4041 different bonding fragments were attached to both nitrogen atoms included in the piperazine core, named Pos1 and Pos2, with no linkers defined between the nitrogen atom and the fragment, where each resulting molecule was considered an independent combination. By combination, it generated a total of 16 329 681 compounds to be screened, an unprecedented number of analyzed ligands compared to gp120 inhibitor studies reported in the literature. To decrease the computational demand required for the minimization of more than 16 million compounds, the next stages of were performed with the gp120 tangled (unminimized) structures.

From the chemical point of view, Combiglide-generated molecules are constituted by the piperazine core and two fragments bonded in different union points, providing new properties (such as solubility, pharmacokinetics, a right number of hydrogen bond donors/acceptors, and molecular weight, to reach the desire biological activity). It is worth mentioning that the initial Glide database only includes fragments derived from small molecules reported in the medicinal chemistry literature, as stated in their website, no larger than 37 atoms and molecular weights below 226 g/mol, making synthetic viability of the generated library considerably higher compared to those randomly generated.

Additionally, a grid box was prepared to perform future docking stages. This box was formed by an innerbox of 10×10×10 Å and an outterbox of 30×30×30 Å, sized for ligands with a length of 20 Å, both centered in residues Trp427, Trp112, Val255, Thr257, Glu370, Ile371, and Asp368 selected (x,y,z coordinates: 27.28, −11.81, 82.61). An OPLS3(16) force field was used to generate the grid box, which included a van der Waals radius scaling factor of 1.0 with a partial charge cutoff of 0.25. A similar grid file is obtained for the gp120 structure coming from the 1G9M PDB file, prepared with the same steps previously explained.

### 2.3 Docking funnel

Molecular docking was performed with Glide(16) taking the generated library through a bottleneck process that narrows the search of drug-like molecules (figure 4). The docking funnel is divided into three steps: High Throughput Virtual Screening quality (HTVS) docking, Standard Precision (SP) quality docking, and Extra Precision (XP) quality docking. To select which ligands were going to remain in the screening analysis, a reference ligand had to be considered. Compound **1** (figure 1) was used as a reference; since a crystallographic structure of this molecule bound inside the Phe43 cavity of gp120 is not available, we carried out an XP quality docking obtaining a score of −7,7 kcal/mol used to filter out all possible hits during the screening. Jobs and advanced settings were set as predetermined by the program, rewarding intramolecular hydrogen bonds and penalizing nonplanar conformation for amides, while planarity of conjugated pi groups was enhanced, and ligand sampling was performed as flexible. Per-residue interaction scores for residues within 12 Å of the grid center were included in the calculation for SP and XP docking screenings

**Figure 4:**
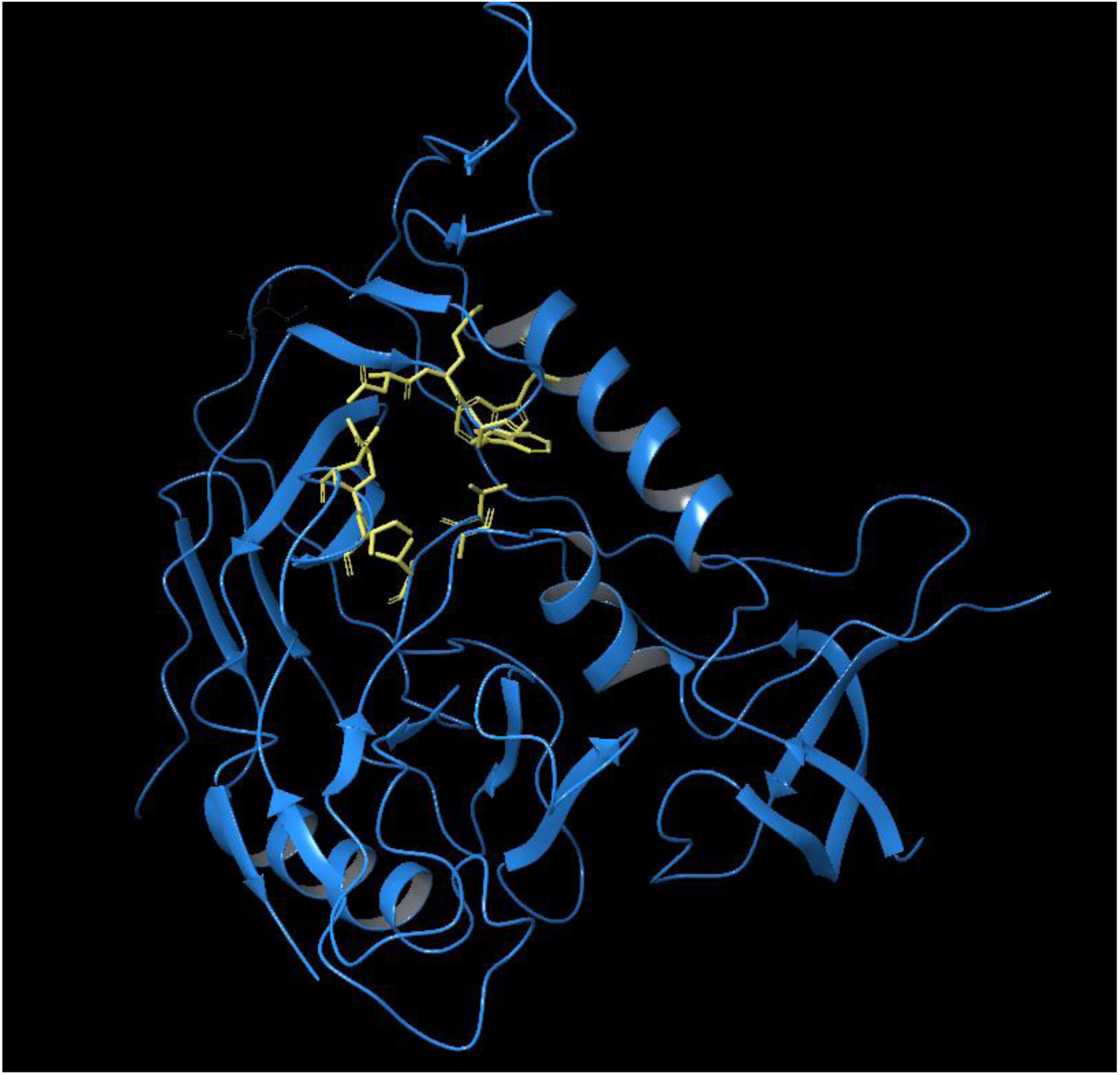
Relevant gp120 residues around its Phe 43 cavity (yellow)

While all the generated ligands were submitted to the first screening, only those ligands within the same score order as the reference ligand (see above) (better than −7.0) were submitted to the SP screening. Nonetheless, ligands submitted to the XP step were previously processed with LigPrep(16) to generate those states that would be found at physiological pH (7.4). Final hits were submitted to consensus docking, as well as to descriptor analysis and biological activities prediction using QikProp(16).

### 2.4 Consensus docking and first ranking

After performing all the screening steps, a consensus docking was carried out to identify those ligands capable of binding with high affinity and getting high docking scores with: a) a structure coming from a different PDB file, and b) a different docking software, as a measure to eliminate biased ligands that would only come as positive candidates with the first gp120 minimized structure, enhancing the possibility to find hits that would bind the protein in any of the conformations presented by the amino acids conforming the Phe43 cavity. 275 hits resulting from the XP step of the docking funnel, named them ligand_001 through ligand_275, and were submitted them to three different calculations: Glide XP quality molecular docking with gp120 from the PDB file 1G9M, AutoDock 4.2.6(16) molecular docking with gp120 from the PDB file 1GC1, and AutoDock molecular docking with gp120 from the PDB file 1G9M

Gp120 envelope glycoprotein was prepared from both the 1GC1 and 1G9M PDB files. The standard preparation workflow was followed, by adding all hydrogens and assigning Gasteiger charges (19,20). Grid maps were calculated following the standard AutoDock protocol for fixed side chains. For the grid box, 55 points were specified in x, y, and z directions, having a spacing factor of 0.375 Å. Extended search parameters were specified in the script. A Lamarckian genetic algorithm was used, with 25 000 000 evaluations, a population size of 150, and 20 runs. A 2.0 Å RMSD tolerance was considered for clustered results.

To obtain the best 100 hits of this stage a ranking function (equation 1) was applied to both Glide XP and AutoDock calculations and their docking score values:

*Equation 1: Consensus docking ranking function*

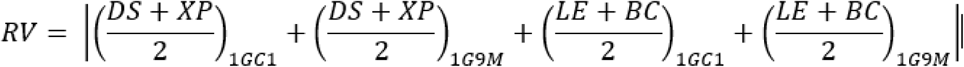

Where the ranking value (RV) is defined by the absolute value of the sum of the docking score (DS) and XP docking score (XP) average, provided by Glide in the two PDB files, 1GC1 and 1G9M (1 and 2, respectively), and the average of the lowest energy (LE) and best cluster (BC) values provided by AutoDock, in both PDB files. Thereafter the ranking function was used to provide a position from 1 to 275 comparing every ligand value against the rest of the molecules included in the set.

### 2.5 Induced-fit docking, second ranking, and visual inspection

Once the top 100 hits had been selected, they were included into the next stage of the project using the Induced-Fit Docking(16) tool from Glide, using the standard protocol with an OPLS3 force field. Side chains within 5.0 Å of ligand poses were refined with Prime, and final structures within 30 kcal/mol of the best structure were redocked with XP quality. The second ranking consisted in the comparison between the absolute values of induced-fit docking score averages by each ligand-receptor complex group of poses (Please refer to supplementary info). These groups were previously filtered by leaving out those ligands with functional groups too reactive or too problematic for future redesign (e.g. esters).

### 2.6 Molecular Dynamics

Simulations were carried out to have a deeper understanding of the ligand-receptor interaction with the help of Desmond(16). These simulations were run during 25 ns in OPLS3 force field, using a TIP4PEW solvent model, an orthorhombic box with minimized volume of 612,144 Å_3_ (distances a=b=c=15Å), chloride ions placement by recalculation (usually 4 or 5 needed to neutralize system), and a 0.15 M NaCl concentration.

## 3. RESULTS AND DISCUSSION

### 3.1 Generation Stage

Despite better performance in clinical trials by compound **3**, in this work we decided to use compound **1** as a scored reference for various reasons: first, as the BMS-378806 prototype, it is an HIV-1 in vitro inhibitor with high potency and low cytotoxicity toward a range of cell lines (6,21), second, its structure is chemically simpler than compounds **2** and **3**, therefore making it easier to synthesize (especially the piperazine moiety, against the chiral methyl-piperazine present in compound **3**), and finally, it yielded a higher docking score, giving us a better cutoff value for the candidate filtering stage explained in the methodology. Finally, inhibitors targeting gp120 are expected to be more effective against HIV infection than those targeting gp41: gp120 binding site to CD4 is exposed in a native state, while gp41 binding sites are exposed after the gp120-CD4 complex is formed (5)

Protein preparation with both PDB files through the protein preparation wizard in Glide, was the first successful step towards generating a grid that would discriminate between ligands fitting the cavity, and those with an inadequate conformation. Ramachandran plots were obtained for both procedures, showing that most calculated angles agreed with the template of energetically allowed regions for backbone dihedral angles in both cases.

Envelope proteins gp120/gp41 have specific affinity with CD4 receptor embedded in CD4+ T cells (22). This interaction is followed by CC or CXC chemokine receptors biding, which are present in the surface of lymphocytes and monocyte/macrophages. These interactions lead to conformational changes in the HIV-1 envelope, eventually ending up in membrane fusion in less than an hour after the first contact between the virus and the T cell.

The deglycosylated core of gp120 is formed by 25 β-strands, 5 α-helices and 10 defined loop segments, also grouped as five conserved constant regions, C1-C5, and five variable loop regions, V1-V5, conforming a prolate ellipsoid with 50×50×25 Å, folded into two major domains (inner and outer) connected through a bridging sheet. Loops in this protein are relatively mobile, and glycosylation sites are all surface-exposed. (5,17)

CD4 co-receptor binds into a depression or cavity formed at the interface between the outer and inner domains, and the bridging sheet. Interaction between both proteins is held by complementary electrostatic potentials: 219 van der Waals interactions and 12 hydrogen bonds are spread around all the contacts between 22 and 26 residues in CD4 and gp120, respectively; among those contacts we can find CD4 residue Phe43 (hence the name of Phe43 cavity) surrounded by the following gp120 residues when inserted into the cavity (figure 4): Trp427, Trp112, Val255, Thr257, Glu370, Ile371, and Asp368, while hydrophobic interactions are found between CD4 Phe43 and gp120 Trp427, Glu370, Gly473, and Ile371, and between CD4 Arg59 and gp120 Val430 (17). Equally important to mention is the fact that residues Trp427, Glu370, and Asp368 are mutational hot-spots, meaning that their substitution would affect CD4 binding, even though Phe43 only interacts with residues Glu370, Ile371, Asn425, Met426, Trp427, Gly73, and Asp368. (17).

At the beginning, minimization of the gp120 protein was performed without any other moieties attached, nevertheless, since the resulting surfaces showed a repositioning of the sidechains of the amino acids, we left CD4 structure bound inside the cavity. When this action was considered, after deleting CD4, Phe43 cavity was clearly present in the structure, eliminating major mistakes at the time of grid generation.

### 3.2 Screening and Validation stage

#### 3.2.1 Reference compound

Compound **1** was docked (figure 5) with extra-precision quality to obtain a reference score value and future cutoff for the library filtering. An XP GlideScore of −7.7 kcal/mol was obtained for this compound with three fragments: a piperazine core, an indole linked through a diacetyl bridge, and a benzamide moiety.

**Figure 5:**
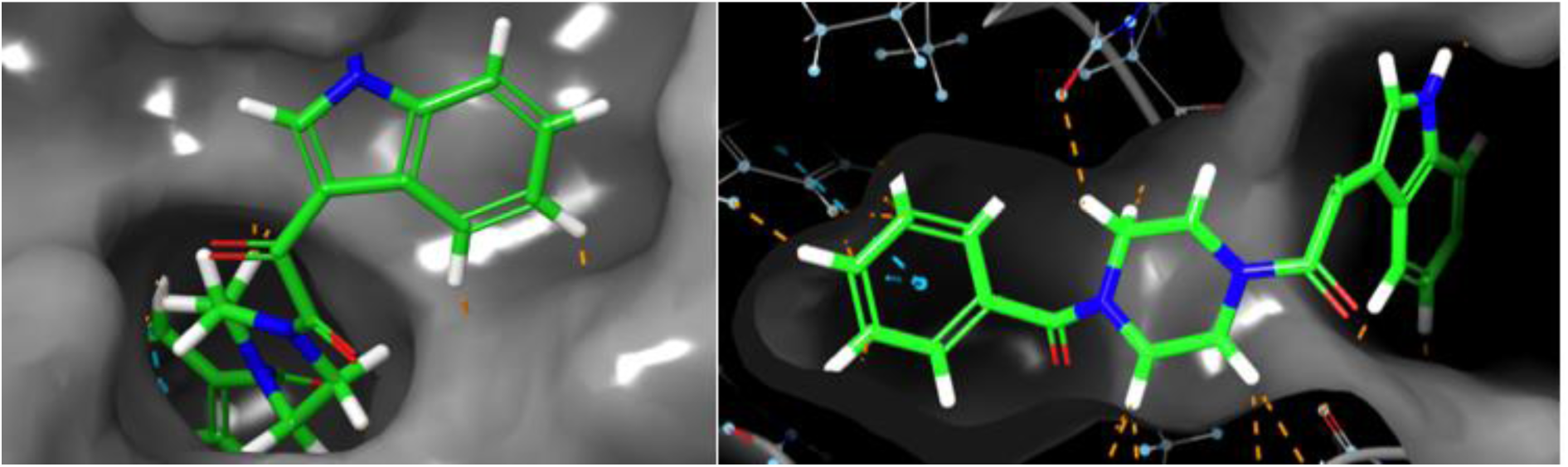
Docked reference compound inside the Phe43 cavity viewed from the outside (left). Complementarity between the cavity and the reference compound. Orange dotted lines indicate hydrogen-bond interactions, while blue ones indicate pi-pi stacking (right).

Compound **1** presented interactions inside and outside the Phe43 cavity (figure 6). The benzamide fragment displayed two pi-pi stacking interactions with Phe382 and Tyr384 residues as the main binding force of the ligand, while the indole moiety was partially exposed to water. The piperazine core occupied the “entrance” of the cavity, surrounded by Trp427, Asn425, Met436, and Gly473 residues, needed to establish an interaction between gp120 and CD4. It is important to bear in mind that interaction diagrams built by Maestro show all residues interacting with the ligand within a 5 Å, but the fact that an amino acid is displayed does not imply that the contact is significant, unless the program indicates so (i.e. H-Bond interactions, ionic interactions, etc.)

**Figure 6:**
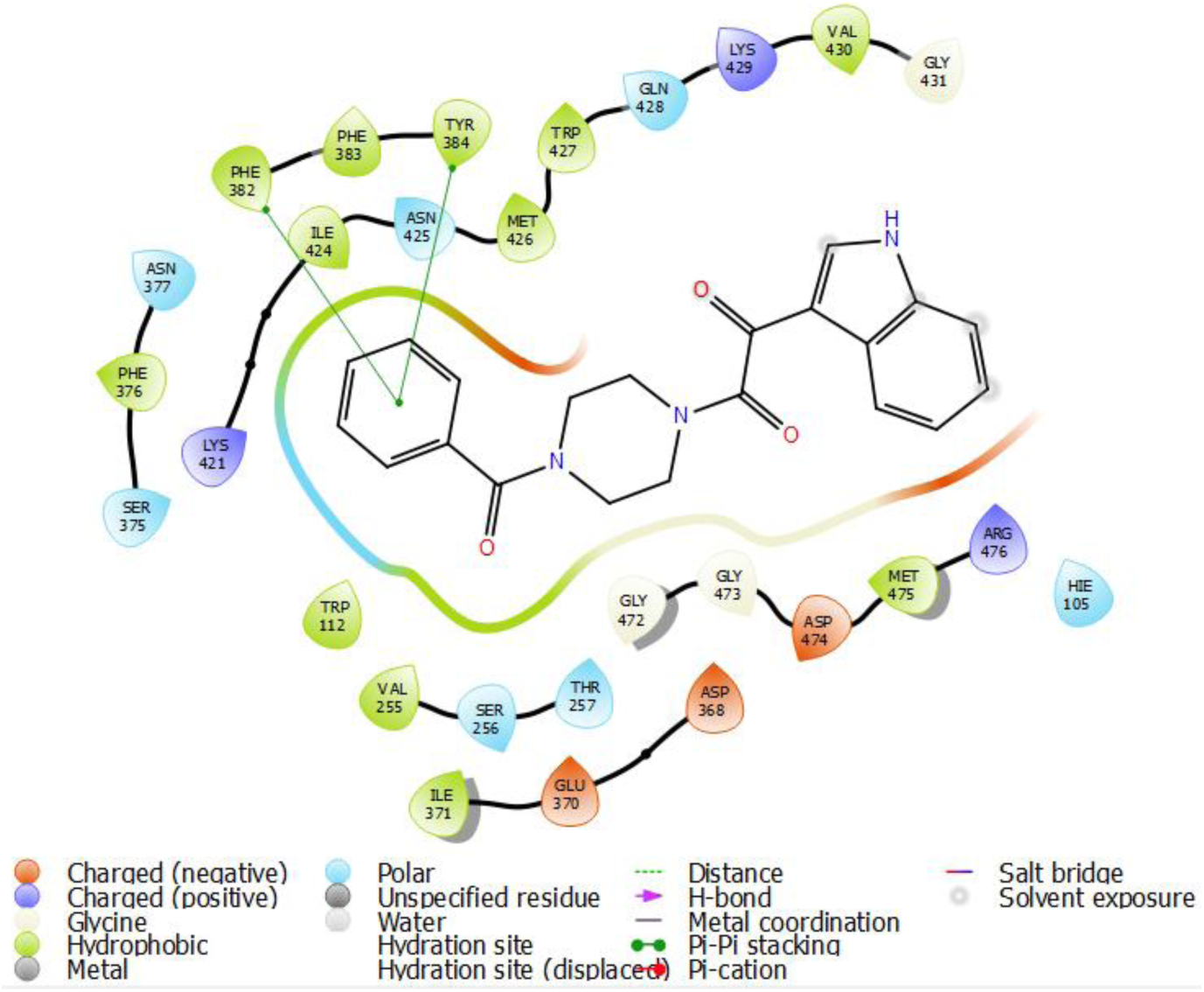
Interactions diagram for the reference compound (compound 1)

The ADME analysis properties are shown in the table 1 (abbreviations can be found in the annex B). Green boxes indicate values inside the recommended thresholds, yellow ones indicate acceptable but not optimal values, while red ones point out undesirable values. Compound **1** shows a majority of optimal values, with exception of the blood/brain barrier coefficient, which explains the −1 value for central nervous system activity.

**Table 1:**
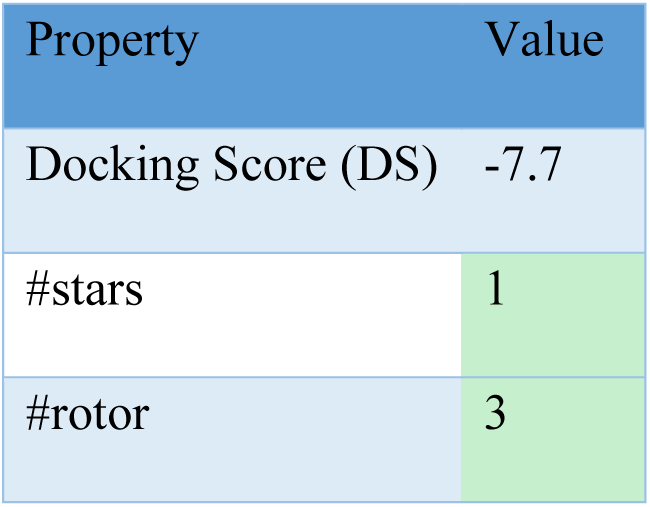
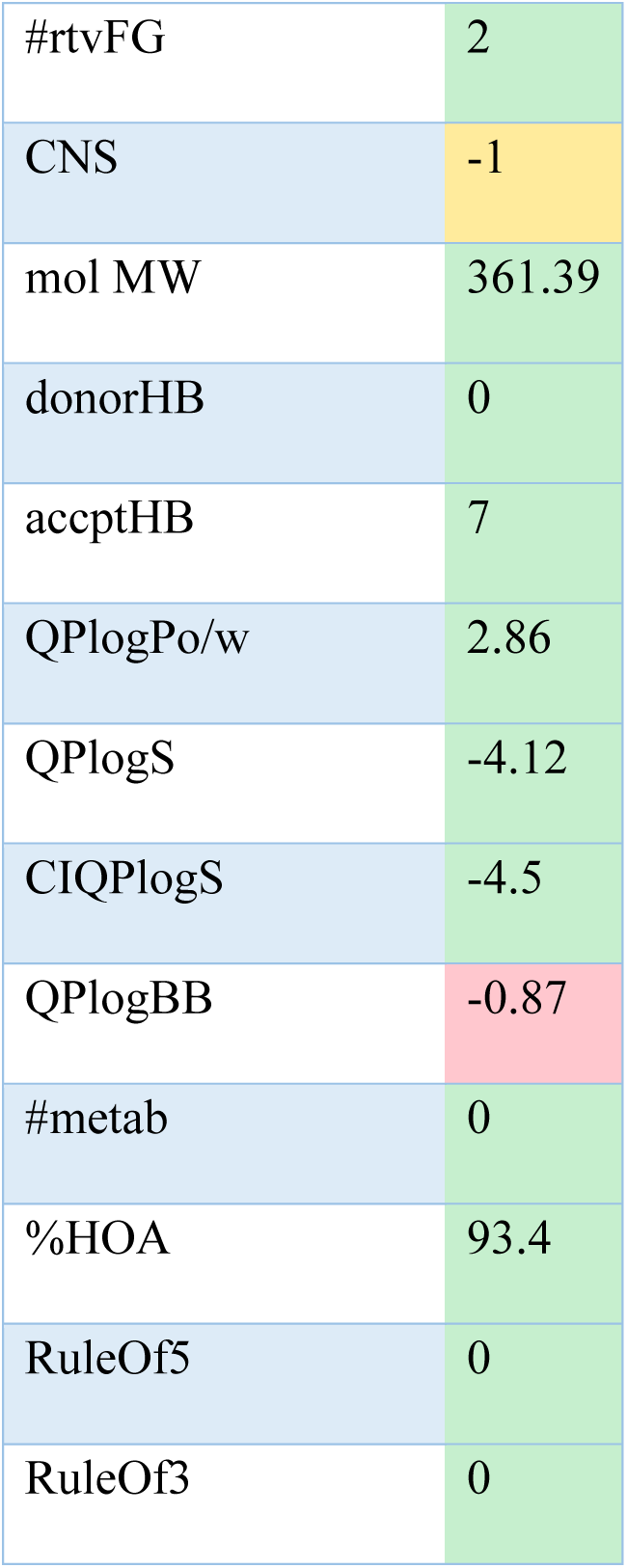
*QikProp properties of reference compound*

| Property | Value |
| --- | --- |
| Docking Score (DS) | -7.7 |
| #stars | 1 |
| #rotor | 3 |
| #rtvFG | 2 |
| CNS | -1 |
| mol MW | 361.39 |
| donorHB | 0 |
| acctpHB | 7 |
| QPlogPo/w | 2.86 |
| QPlogS | -4.12 |
| CIQPlogS | -4.5 |
| QPlogBB | -0.87 |
| #metab | 0 |
| %HOA | 93.4 |
| RuleOf5 | 0 |
| RuleOf3 | 0 |

#### 3.2.2 Docking Funnel

As described in the methodology, high-throughput virtual screening was the start of the filtering process for a total of 16 329 681 ligands. The docking score cutoff value used for this stage was −7.0 kcal/mol, to include all those molecules within the same affinity range or order of magnitude than the reference compound (−7.7 kcal/mol), leaving 60,379 molecules to work with during the next step in the docking funnel.

Molecules with lower (better) values than the cutoff represent around 0.37% of the initial size of the library. This means that the majority of compounds are not suitable for gp120 inhibition because either the compound scores poorly and lacks stable interactions, or the size and conformation of the molecule prevent it from fitting inside the cavity, in which case the program does not score it.

While the HTVS docking stage served primarily as a way of eliminating all ligands with low scores, leaving those worth working with, the SP docking stage reinforced the evaluation of the remaining portion of the library through a more detailed scoring function. For this stage, a cutoff value of −7.7 kcal/mol was used to refine those compounds with equal or better activity than the reference, leaving a total of 12,446 ligands, or nearly 21% of the initial ligands for this step.

Resulting ligands of the SP docking stage were submitted to the XP docking stage after they underwent a conformational analysis with LigPrep. From the initial 12,446 ligands, a final number of 275 compounds (2.2%) was obtained with a cutoff value of −7.7 kcal/mol and the most robust scoring algorithm available within Glide.

It is equally important to mention that scores through the docking funnel ranged between −1.2 and −9.7 kcal/mol, with obvious exception of ligands that did not fit into the cavity. According to the equation Δ*G* = −*nRTlnK*_*af f*_, where Δ*G* represents the Gibbs free energy change for the formation of the protein-ligand complex and *K*_*af f*_ stands for the affinity constant of the same complex, we can obtain the difference of affinity between a ligand and the reference through the following expression: 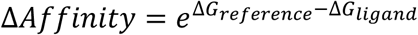. This means that the screened library contained ligands with affinities 7.4 times higher than the reference.

#### 3.2.3 Consensus docking and first ranking

Once the docking funnel obtained the best 275 hits out of the initial library, the ligands were renamed 1 through 275 for future calculations. Consensus docking was performed using the 1G9M PDB file, and the AutoDock 4.2.6 software to eliminate those ligands binding only the gp120 structure created by minimization of the 1GC1 PDB file. This step becomes necessary when trying to separate those molecules biased to bind only one specific conformation of the protein from those that truly establish interactions with significant amino acids. Still, this is only helpful when both structures are structurally equal or very similar; for example, 1G9M and 1GC1 differ only by a mutation, which is not located in the region of our interest.

Even though scoring functions for both programs are different, a similarity in the order of the results is noticeable among them and both PDB files. The ranking shows in green the best 100 compounds, in yellow the following 100 best ligands, and in red the last 75 ones. Nevertheless, these compounds can still be analyzed and redesigned to become bioactive compounds, since they still figure as the best virtual hits from this library.

### 3.3 Refinement stage

After the screening and validation stage was finished, best ligands needed to be studied in more detailed through the refinement stage, where induced-fit docking (IFD) and molecular dynamics gave us a better insight of the calculated interactions.

#### 3.3.1 Induced-Fit docking, second ranking, and visual inspection

After the first ranking was performed, the 100 best ligands were submitted to IFD calculations to find the best possible poses of the ligand inside the Phe43 cavity, and the best conformations of the protein when in contact with the ligand. After output poses where averaged and ranked as a simple comparison of the absolute values of the IFD score, the best 20 compounds were obtained.

These 20 compounds were pre-filtered before resulting the best ligands. None of these compounds are expected to present undesired reactivity under physiological conditions and their metabolic degradation should proceed without further consequences for the organism. This represents two main advantages: one, the compounds are expected to be stable when isolated as final synthesis products, and two, they are less likely to present unusual reactivity or side reactions when tested in biochemical assays.

The reader must be aware that the generated library is made of symmetrical compounds, where Pos1 and Pos2 of the attached fragments can be the same in two different ligands, being the same compound. Looking at every ligand in the table, we noticed that 3 pairs of ligands were the same compounds, with ligand 140 being the same one as ligand 141, ligand 73 the same as ligand 74, and ligand 59 the same as 155. This shows that the methodology in this work is reproducible within the same set of molecules, bringing those ones that are similar between them into positions relatively close, despite the consensus docking stage.

##### 3.3.1.1 Top Ten Ligands

###### Ligand 255

Ligand 255 (figures 8-10) presented three interactions in docking calculations: An H-bond between the nitrogen atom contained in the 4,5,6,7-tetrahydroindole fragment and Asn425 residue, a hydrophobic interaction between the same fragment and Trp427 residue, and a H-bond between the amino group contained in the phenylsulfonamide moiety and the Trp427 residue. All the residues in classes A and B are present around this ligand, and exposure to water is only displayed for the oxygen atom of the sulfonamide group. On the other hand, the MD studies show complete exposure to water during the simulation, possibly due to variations in the size of the cavity that allowed water molecules to move inside it during the first nanoseconds. Two of the most important interactions detected in docking calculations, Trp 427 and Asn 425, were confirmed, occurring during 88% and 93% of the simulation time, respectively. A new interaction, an H-bond between the sulfonamide and the Met 475, was proven relevant with 74% of the simulation time.

**Figure 7:**
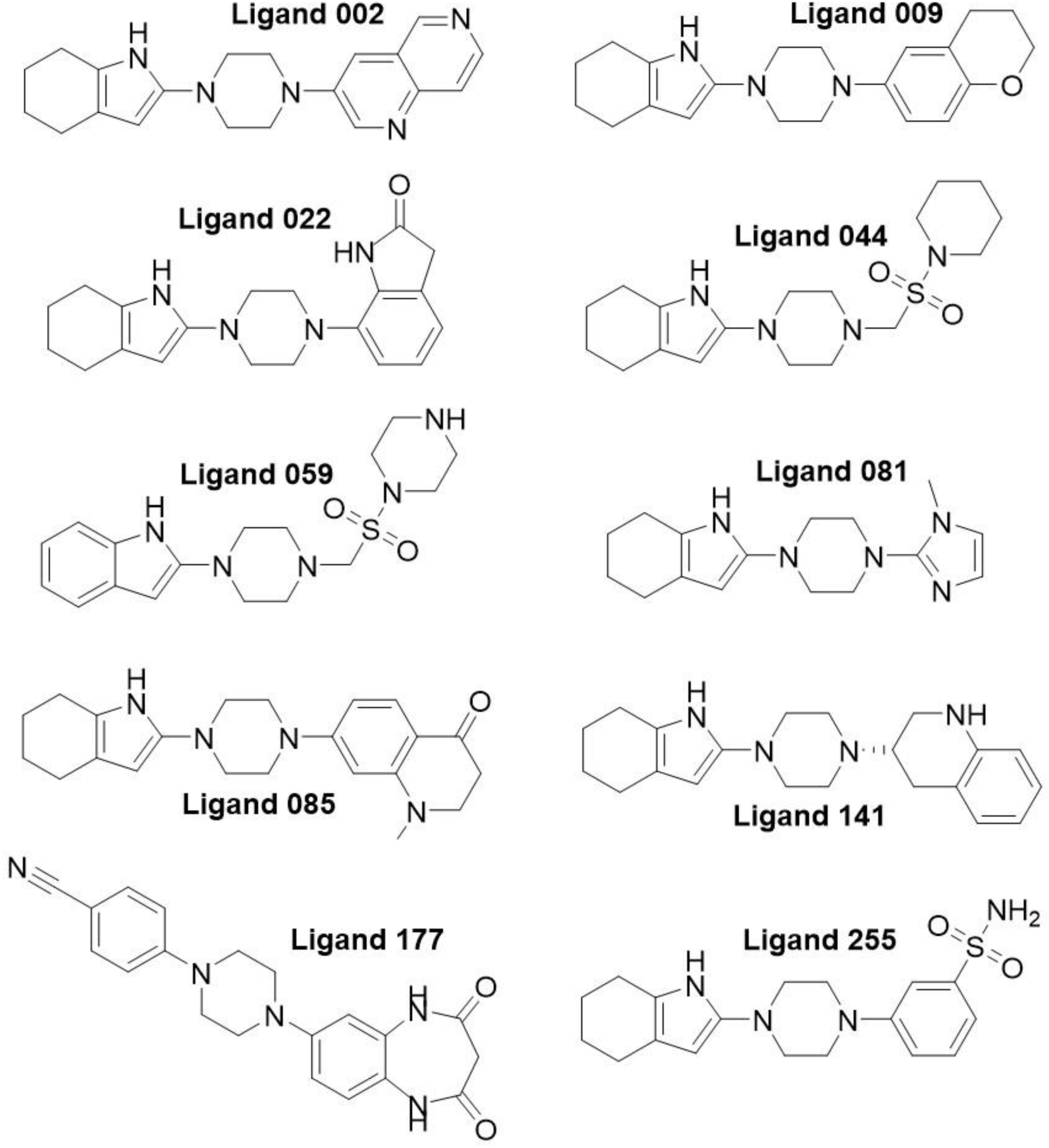
Top-ten virtual hits obtained at the end of the docking funnel

**Figure 8:**
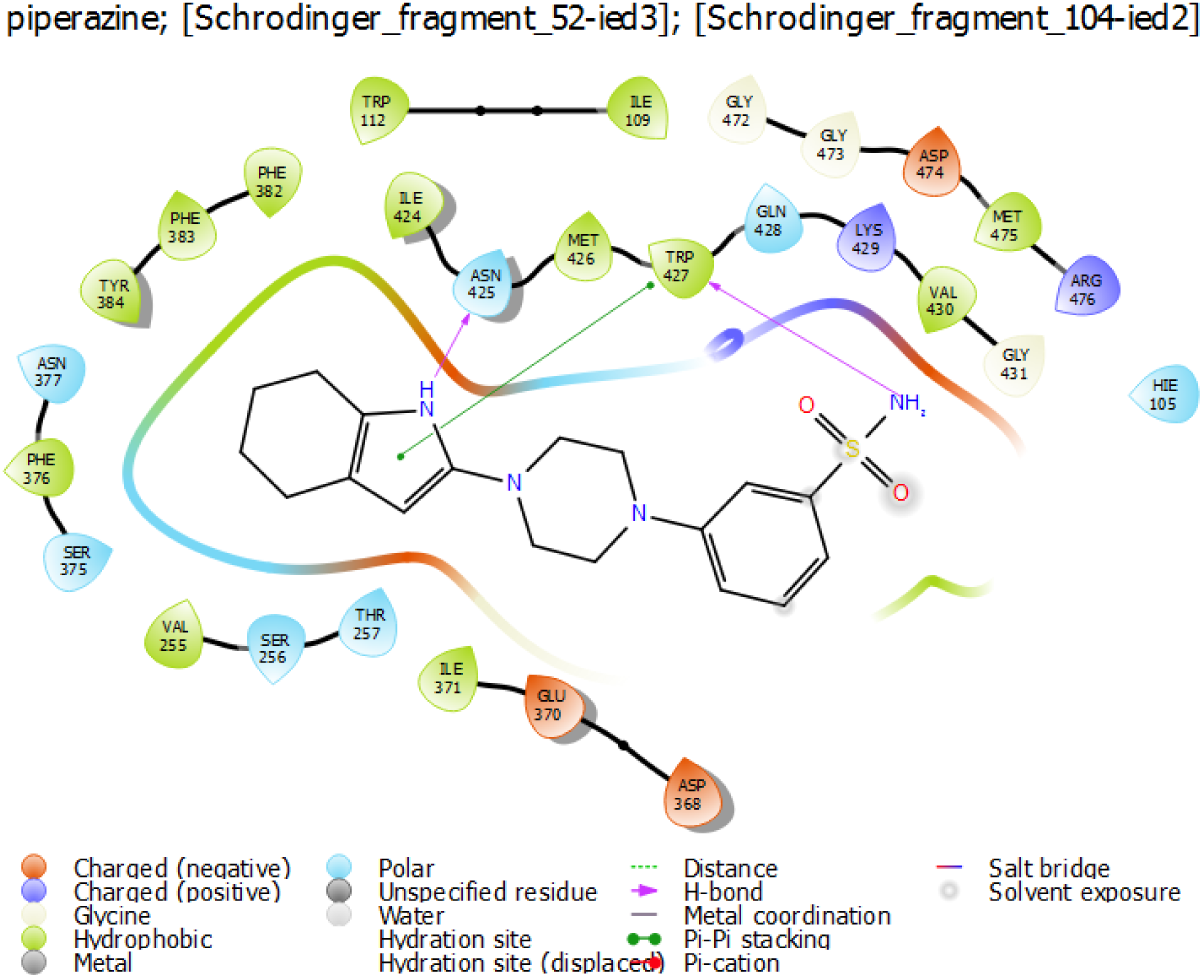
*IFD interaction diagram for ligand 255*

**Figure 9:**
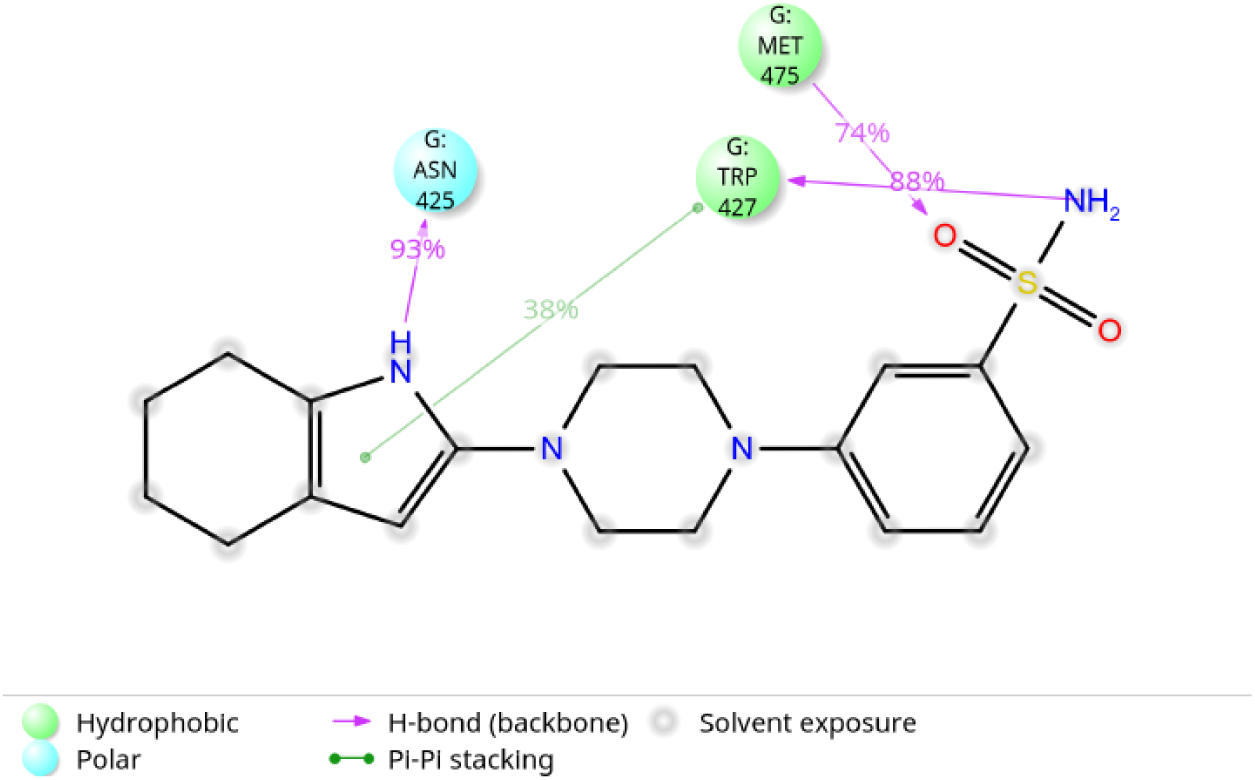
Ligand 255 (MD1) simulation interactions diagram

**Figure 10:**
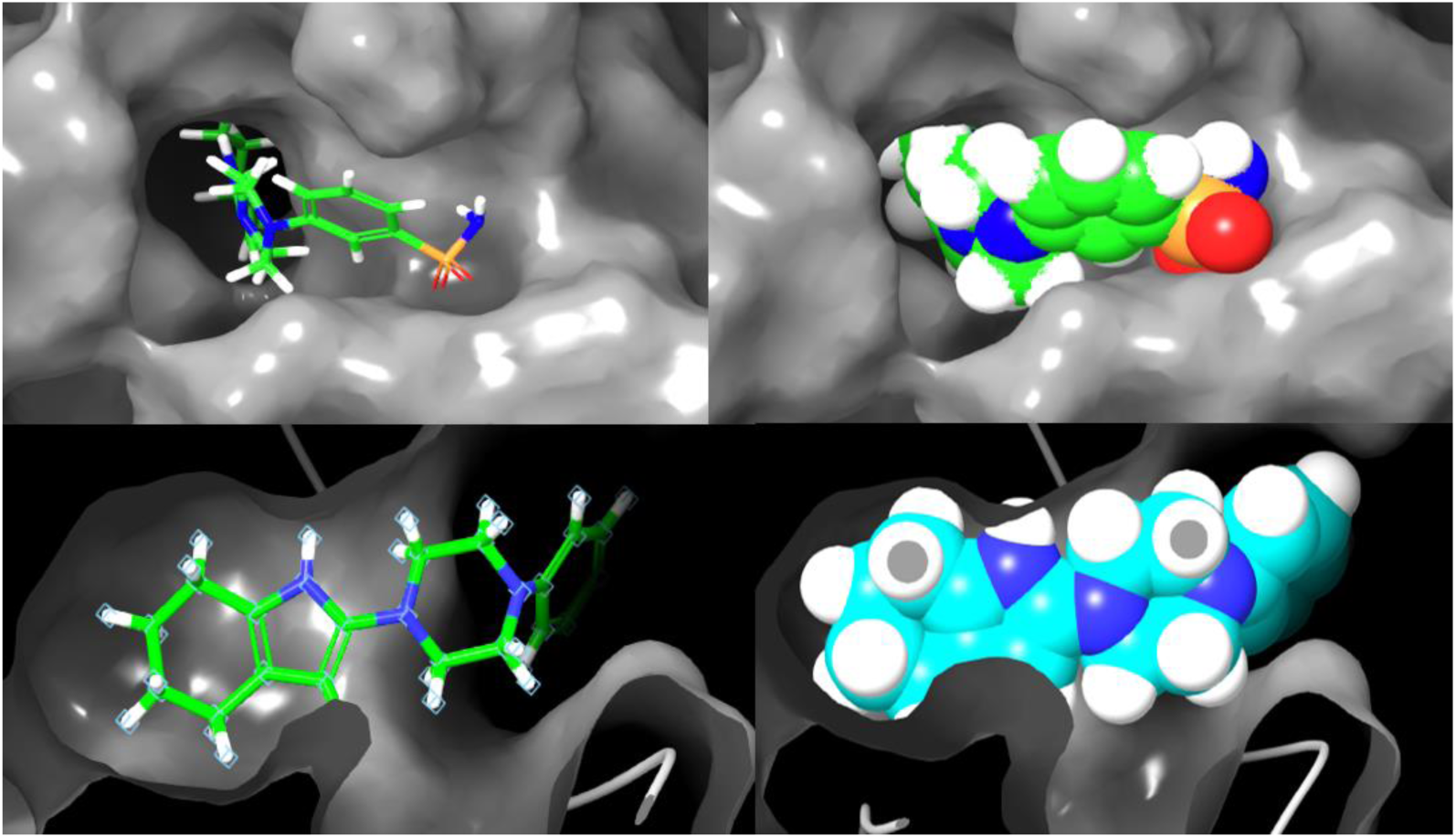
Ligand 255 from different points of view, inside the Phe 43 cavity in gp120. Its complementarity with the pocket, and the position of the piperazine ring at the cavity entry are noticeable at first glance.

###### Ligand 141

Four main interactions were displayed for the docking calculations of ligand 141: hydrophobic contact between the Trp 427 residue, H-bond between the nitrogen atom of the 4,5,6,7-tetrahydroindole fragment and the Asn 425 residue, and two H-bonds between the protonated nitrogen atom in the piperazine core and the Asp 368 residue (all class B interactions). All residues from classes A and B were present within 5 Å of the ligand contact zone. Water exposure was only displayed for the 1,2,3,4-tetrahydroquinoline fragment lying outside the Phe 43 cavity.

Unlike ligand 255, ligand 141 simulation confirmed both the interactions and the level of water exposure. Interactions for this ligand were present most of the simulation time. For the Asp 368 interaction, it was revealed that a small fraction of it corresponded to ionic contact between the residue and the positive charge of the amino group. This last piece of information, however, is subject to experimental evidence, since the nitrogen atom is not the only one that can be protonated under physiological conditions.

###### Ligand 002

Ligand 002 docking calculations only showed one strong interaction between the nitrogen atom owned by the 4,5,6,7-tetrahydroindole fragment and Asn 425, confirmed as a 98% simulation time contact by the MD studies, which also revealed a purely hydrophobic interaction between this fragment and Trp 427 amino acid. Classes A and B interactions were also present within the contact volume. While docking calculations showed solvent exposure for the [1,6]-naphtyridine fragment alone, MD studies displayed it for the entire ligand during simulation time, possibly for the same reasons as ligand 255.

###### Ligand 081

Ligand 081 showed the same interactions as ligands 255, 141, and 002, regarding the 4,5,6,7-tetrahydroindole moiety. The main differences between ligand 081 and other top 10 molecules from this library, were pointed out by the MD studies: First, the Trp 427 hydrophobic interaction is particularly low in the percentage of simulation time, with only 39%, second, Phe 376 residue maintains an interaction (lower than 30% simulation time) with the same fragment at the bottom of the Phe 43 cavity without being neither a class A nor a class B interaction, and most important, docking and MD studies differ by an amino acid interacting through an H-bond with the 1-methylimidazole moiety, Gly 473, and Asp 368.

###### Ligand 009

This virtual hit presents almost the same interactions as ligand 081, with slightly better simulation times for residues Trp 427 and Asn 425. Despite durable bonds in the inner area of the cavity, the chromane moiety in the exterior does not present any interactions that might anchor the ligand outside the hydrophobic pocket, explaining the total solvent exposure shown in the simulation diagram.

###### Ligand 022

This ligand shows typical interactions for the 4,5,6,7-tetrahydroindole fragment, displaying a low 37% simulation time hydrophobic interaction with Trp 427 that is not predicted by docking. At the same time, the oxindole fragment presents several contacts of great interest. Docking studies show this fragment interacting through a donor H-bond between the nitrogen atom and the Gly 473 residue, and acceptor H-bond between the carbonyl group and the Met 475 residue, an amino acid that is not included in any of the interaction classes described at the beginning of this section.

On the other hand, MD simulations show no interaction with the Gly 473 residue, instead displaying a water-mediated double interaction between residues Asp 474 and Met 475, and the carbonyl group. Crystallization of the complex would be needed to confirm if this water molecule is required to establish an interaction in the outer side of the pocket, or if it happens to just be a statistically significant interaction throughout the simulation with different water molecules.

###### Ligand 059

Ligand 059 showed a total of 5 interactions in docking calculations: a double hydrophobic interaction with Trp 427 from both rings forming the indole fragment, an H-bond from the nitrogen atom of this fragment towards Asn 425, and two H-bonds coming from the protonated nitrogen atom present in the 1-(methylenesulfonyl)piperazine with Asp 368. All of these class B interactions were confirmed by MD studies with simulation times higher than 90%, except for Trp 427 contacts. Solvent exposure was also shown to be the same for both studies.

###### Ligand 177

Along with ligand 059, ligand 177 has a different first fragment than the other virtual hits. The benzonitrile fragment presented no interactions within the inner section of the Phe 43 pocket in docking calculations, even though this cavity is highly hydrophobic. Nevertheless, poor interactions between this fragment and Trp 427 and Met 475 residues was documented by MD simulation, leading to fragile anchorage of the ligand inside the gp120 protein.

On the other hand, the 1H-1,5-benzodiazepine-2,4(3H,5H)dione moiety showed multiple interactions, albeit different between both analyses. Docking showed a double accepting H-bond between residues Arg 475 and His 105 (none of them class A or B) and one of the carbonyl groups, and a donor H-bond towards Trp 427 coming from one of the nitrogen atoms of the benzodiazepine group, while MD studies displayed one of these nitrogen atoms in contact with Met 426 residue through a poor 32% H bond, and its second nitrogen forming an occasional water-bridged contact with Asp 368. As it happened with previous hits, this molecule showed total solvent exposure throughout the simulation.

###### Ligand 085

Even though class A and B interactions were present within 5 Å from the molecule, only the 4,5,6,7-tetrahydroindole fragment presented interactions like the ones described in previous ligands, as well as full solvent exposure inside the pocket. From both, docking and MD studies, we determined that the 1-methyl-2,3-dihydro-4(1H)-quinolinone moiety did not present significant contacts that might improve the ligand’s inhibitory capacity.

###### Ligand 044

Structurally and conformationally similar to ligand 059, ligand 044 presented typical interactions for the 4,5,6,7-tetrahydroindole moiety in both analyses. Solvent exposure was shown only for the piperidine group present in the 1-(methylenesulfonyl)piperidine fragment. This last group showed no interactions in docking calculations, but one H-bond contact through a bridging water molecule with Asp 368 in the MD simulation.

##### 3.3.1.2 Molecular dynamics quality parameters

As a first measure to monitor simulation quality, RMSD of the alpha carbon atoms of the protein complexes was recorded during the simulations, and then compared to that RMSD of the protein simulated on its own.

All of the simulations converged within 5 ns, a relatively small amount of time for medium-size systems such as gp120, having an RMSD of less than 5 Å during the following 20 ns. The exceptions to this description were simulations MD3 and MD5, that converged around 7-8 Å, and 5-6 Å respectively.

Alternatively, the protein RMSF was also recorded for each one of the simulations. The clear majority of residues displayed an RMSF value between 1-2 Å, showing low to zero fluctuation for most amino acids contained in gp120. However, it is worth discussing those peaks rising above 2 Å:

1. **Residues 120-140:** Showing the highest RMSF values, up to 15 Å, these residues belong to the V1/V2 loop and β2 region of gp120 protein, the most mobile region in the protein when detached from the whole viral structure. Nonetheless, this area is also in contact with any ligand inside the cavity, making it susceptible to fluctuations due to existing contacts, clashes, and interactions.
2. **Residues 150-160:** As a part of the V1/V2 loop, their 3 Å fluctuations might be caused by the loop’s natural movement.
3. **Residues 270-280:** These residues with 3 Å fluctuations belong to the β10 region, usually affected by movements made by domain V3 loop, next to it.
4. **Residues 300-320:** The second most mobile region of the protein with RMSF values of up to 6 Å, these amino acids form the V3 loop, a region responsible for chemokine attachment once gp120-CD4 interaction has been achieved, triggering conformational changes.
5. **Residues 340-350:** With values of 3 Å, these residues belong to the α2 helix of gp120. These changes cannot be attributed to the effect of the ligand, taking into consideration the distance between the helix and the subdomains surrounding the Phe 43 pocket.
6. **Residues 365-370:** This region has a peak RMSF value of 4 Å, and belongs to the β15 strand. The fluctuations presented in this region are caused by the presence of the ligand, since this strand interacts with the C‘’ strand of CD4.

#### 3.3.2 Biological properties prediction (ADME)

A set of endpoints and descriptors were calculated with three programs (QikProp, T.E.S.T., and Derek Nexus)(16) to study the predicted chemical and biological properties of the best virtual hits. Table 2 shows the 15 descriptors calculated by QikProp according to the recommended range of values in each one. Values in green are ideal, values in yellow are acceptable, and values in red represent properties out of the recommended threshold. While all compounds had better docking scores than the reference, not all properties were ideal for these hits. Blood-brain barrier partition coefficient of most compounds, except ligands 177 and 255, were not acceptable, and most important, lower than the reference compound. This descriptor is directly related to the activity in central nervous system, for which ligands 177 and 255 presented better values than the reference. Finally, a couple of compounds presented predicted human oral absorption values below 80%, for which experimental tests would need to be developed.

**Table 2:** *QikProp descriptors comparison between reference compound and top 10 ligands*

| Ligand | Docking score | #stars | #rotor | #rtvFG | CNS | mol MW | donorHB | acceptHB | QPlogPo/w | QPlogS | CIQlogS | QPlogBB | #metab | %HOA | RuleOffive | RuleOTThree |
| --- | --- | --- | --- | --- | --- | --- | --- | --- | --- | --- | --- | --- | --- | --- | --- | --- |
| 002 | -8.993 | 0 | 0 | 0 | 0 | 333.44 | 1 | 4.00 | 4.38 | -6.37 | -5.37 | -0.08 | 6 | 100 | 0 | 1 |
| 009 | -8.826 | 3 | 0 | 0 | 1 | 337.46 | 1 | 2.25 | 5.51 | -7.20 | -5.94 | 0.41 | 4 | 100 | 1 | 1 |
| 022 | -8.599 | 0 | 0 | 0 | 0 | 336.44 | 2 | 4.00 | 3.95 | -6.20 | -5.22 | -0.28 | 4 | 100 | 0 | 1 |
| 044 | -8.322 | 0 | 3 | 0 | 1 | 366.52 | 1 | 7.00 | 2.66 | -4.11 | -3.24 | -0.12 | 4 | 87.964 | 0 | 0 |
| 059 | -8.24 | 0 | 3 | 0 | 1 | 363.48 | 2 | 8.50 | 1.01 | -1.67 | -1.89 | 0.03 | 1 | 62.015 | 0 | 0 |
| 081 | -8.157 | 1 | 0 | 0 | 1 | 285.39 | 1 | 2.50 | 4.38 | -5.82 | -4.66 | 0.28 | 3 | 100 | 0 | 1 |
| 085 | -8.143 | 2 | 0 | 0 | 0 | 364.49 | 1 | 4.50 | 4.68 | -7.07 | -5.67 | -0.13 | 6 | 100 | 0 | 1 |
| 141 | -7.973 | 1 | 0 | 0 | 2 | 336.48 | 2 | 3.50 | 4.24 | -5.32 | -4.28 | 0.64 | 6 | 100 | 0 | 0 |
| 177 | -7.891 | 0 | 1 | 0 | -2 | 361.40 | 2 | 8.50 | 1.45 | -6.12 | -4.96 | -1.93 | 2 | 67.512 | 0 | 1 |
| 255 | -7.729 | 0 | 2 | 0 | -2 | 360.47 | 3 | 6.00 | 2.66 | -5.46 | -4.78 | -1.16 | 3 | 87.393 | 0 | 0 |
| Reference | -7.7 | 1 | 3 | 2 | -1 | 361.39 | 0 | 7.00 | 2.86 | -4.12 | -4.50 | -0.87 | 0 | 93.4 | 0 | 0 |

Table 3 shows two endpoints calculated with T.E.S.T., oral LD50 in rats and mutagenicity (Ames test). No ligands, except for ligands 044 and 009, showed toxicities below 500 mg/*K*g, while seven of them presented no evidence of mutagenic behavior. Ligand 255 again showed improved characteristics against the other 9 virtual hits, with an LD50 around 900 mg/*K*g (Class III of toxicity: slightly toxic), and mutagenicity negative. Results showed in this table next to an asterisk were obtained through nearest neighbor method, which might be taken with caution, even if it is a valid approach. Results expressed as N/A mean a predictive quantitative structure-activity relationship model could not be generated for this molecule.

**Table 3:** T.E.S.T values for LD50 and mutagenicity.

| Ligand | Endpoints |  |
| --- | --- | --- |
|  | LD50 oral<br>rat mg/Kg | Mutagenicity<br>(Ames) |
| Ligand 002 | 609.72 | Negative |
| Ligand 009 | 245.77 | Negative |
| Ligand 022 | 569.14 | Negative |
| Ligand 044 | 289.65 | Negative |
| **Ligand 059** | 505.84* | Positive* |
| **Ligand 081** | 666.06* | Negative* |
| **Ligand 085** | 531.35 | Negative |
| **Ligand 141** | 1144.61* | N/A |
| **Ligand 177** | 811.17 | Positive |
| **Ligand 255** | 908.74 | Negative |

Finally, 60 endpoints where calculated with the help of Derek Nexus in several species. Table 4 shows the results for human, monkey and rat predictions of at least an “Equivocal” level alert. The clear majority of endpoints were not fired (positive activity) for the studied ligands, except for the ligands indicated for each species as Equivocal (EQ) or Plausible (PL). No ligands fired endpoints on Probable or Certain levels, showing a promising profile for future development and minimization of negative consequences in clinical trials. Focusing on ligand 255, all endpoints fired are due to the presence of the sulfonamide group; this could be redesigned to minimize de possibility of a negative biological effect, while maintaining its overall safe profile as an inhibitor.

**Table 4:** Fired endpoints in human, monkey and rat for top-ten ligands in Derek Nexus

| Endpoint | Human | Monkey | Rat |
| --- | --- | --- | --- |
| **5alpha-Reductase inhibition** | None | None | None |
| **Adrenal gland toxicity** | None | None | None |
| **alpha-2-mu-Globulin nephropathy** | None | None | None |
| **Anaphylaxis** | None | None | None |
| **Androgen receptor modulation** | None | None | None |
| **Bladder disorders** | None | None | None |
| **Bladder urothelial hyperplasia** | 255 (EQ) | 255 (PL) | 255 (PL) |
| **Blood in urine** | None | None | None |
| **Bone marrow toxicity** | None | None | None |
| **Bradycardia** | None | None | None |
| **Carcinogenicity** | None | None | None |
| **Cardiotoxicity** | None | None | None |
| **Cerebral oedema** | None | None | None |
| **Chloracne** | None | None | None |
| **Cholinesterase inhibition** | None | None | None |
| **Chromosome damage in vitro** | None | None | None |
| **Chromosome damage in vivo** | None | None | None |
| **Cumulative effect on white cell count and immunology** | None | None | None |
| **Cyanide-type effects** | None | None | None |
| **Developmental toxicity** | None | None | None |
| **Glucocorticoid receptor agonism** | None | None | None |
| **Hepatotoxicity** | 255 (EQ) | 255 (EQ) | 255 (PL) |
| **HERG channel inhibition in vitro** | 059 and 141 (PL) | 059 and 141 (PL) | 059 and 141 (PL) |
| **High acute toxicity** | None | None | None |
| **Irritation (of the eye)** | None | None | None |
| **Irritation (of the gastrointestinal tract)** | None | None | None |
| **Irritation (of the respiratory tract)** | None | None | None |
| **Irritation (of the skin)** | None | None | None |
| **Kidney disorders** | None | None | None |
| **Kidney function-related toxicity** | None | None | None |
| **Lachrymation** | None | None | None |
| **Methaemoglobinaemia** | None | None | None |
| **Mitochondrial dysfunction** | None | None | None |
| **Mutagenicity in vitro** | None | None | None |
| **Mutagenicity in vivo** | None | None | None |
| **Nephrotoxicity** | 177 (EQ), 255 (PL) | 177 (EQ), 255 (PL) | 177 (EQ), 255 (PL) |
| **Neurotoxicity** | None | None | None |
| **Non-specific genotoxicity in vitro** | None | None | None |
| **Non-specific genotoxicity in vivo** | None | None | None |
| **Occupational asthma** | None | None | None |
| **Ocular toxicity** | 255 (PL) | 255 (PL) | 255 (PL) |
| **Oestrogen receptor modulation** | None | None | None |
| **Oestrogenicity** | None | None | None |
| **Peroxisome proliferation** | None | None | None |
| **Phospholipidosis** | None | None | None |
| **Photo-induced chromosome damage in vitro** | None | None | None |
| **Photo-induced non-specific genotoxicity in vitro** | None | None | None |
| **Photo-induced non-specific genotoxicity in vivo** | None | None | None |
| **Photoallergenicity** | None | None | None |
| **Photocarcinogenicity** | None | None | None |
| **Photomutagenicity in vitro** | None | None | None |
| **Phototoxicity** | 255 (PL) | 255 (PL) | 255 (PL) |
| **Pulmonary toxicity** | None | None | None |
| **Respiratory sensitisation** | None | None | None |
| **Splenotoxicity** | None | None | None |
| **Teratogenicity** | 177 (EQ) | 177 (EQ) | 177 (EQ) |
| **Testicular toxicity** | None | None | None |
| **Thyroid toxicity** | None | None | None |

| Uncoupler of oxidative phosphorylation | None | None | None |
| --- | --- | --- | --- |
| Urolithiasis | 255 (PL) | 255 (PL) | 255 (PL) |

## 4. CONCLUSIONS

The bottleneck methodology applied in this work allowed us to explore a much bigger chemical space than those ones reported at experimental studies, with a lower use of resources and time. Descriptors used in this work were sufficient to predict chemical structures with affinities up to 3 times higher, for the gp120 protein, than those reported as active in preclinical studies. At the same time, with this methodology we narrowed down a 16.3 million compounds database to a few molecules that can be analyzed, synthesized, assayed, and redesigned afterwards, within a short amount of time, and less resources.

Thanks to the analysis of all interactions established between gp120 and the virtual hits, it becomes possible to point out at a first scaffold to inhibit CD4 attachment: a piperazine core with a 4,5,6,7-tetrahydroindole moiety as a first fragment interacting in the inner section of the cavity, and a second fragment capable of forming H-bond/ionic interactions in the external region of the cavity. Even though the piperazine did not present any interactions with the surroundings, its conformation and volume are key for blocking the Phe43 cavity. Most important residues include Asn425 and Trp427, key amino acids in CD4 Phe43 interaction. These residues were present in almost all the interactions of our virtual hits, giving us a hint to redesign and improve functional groups in contact with them, in order to achieve better simulation time percentages.

Molecular dynamics studies confirmed the validity of the results obtained by the docking stages, since the ligand interactions were present towards the same amino acids in the majority of cases, and the quality parameters were good throughout the complete trajectories, whose 25 ns simulation time was much longer than other in silico studies reported in the literature. This last point was proved through quick convergence of the RMSD values (5 ns) and low deviations (5 Å).

Simulations showed that these ligands do not get out of the Phe 43 pocket, due to strong and durable H-bond interactions, and despite the solvent exposure of more than half the hits inside the cavity. RMSF values demonstrated that ligands have a significant effect on the V1/V2 loop region, and the β15 strand. These could lead to important conformational fluctuations that prevent gp120-CD4 interaction.

Biological properties prediction displayed great profiles for many of the top-ten ligands discussed in this work. While ligand 255 showed the most promising results on these characteristics, the rest of the molecules could also prove potential candidates during clinical trials, subject to redesign to improve inhibitory activity.

Novelty of this work resides in the combinatorial and massive generation of libraries of compounds that would possibly inhibit gp120-CD4 interaction by insertion inside the Phe43 cavity. The docking funnel described in the methodology represents a powerful and robust tool to discriminate between ligands with lower or higher binding affinity. Finally, we hope that the description of the methodology made in this work will help perform high throughput virtual screening of large libraries with different biological targets and diseases. Future directions include the improvement of the methodology, the testing of new cores and generation of new libraries, as well as the reduction of computational resources needed to perform the screening.

## ACKNOWLEDGEMENTS

We thank the General Direction of Computing and Information and Communication Technologies of the National Autonomous University of Mexico (DGTIC UNAM), for the access to the supercomputer cluster *Miztli*. DJCP also thanks the Institute of Chemistry UNAM, the Council of Science and Technology of the State of Mexico (COMECYT), and the UAEMex foundation for the funding granted. We also thank Francisco Casañas and Citlalit Martínez for the IT support provided.

## ABBREVIATIONS

HTVS: high-throughput virtual screening
HIV: human immunodeficiency virus
AIDS: acquired immunodeficiency syndrome
NRTIs: nucleoside reverse transcriptase inhibitors
NNRTIs: non-nucleoside reverse transcriptase inhibitors
PIs: protease inhibitors
ADME: absorption, distribution, metabolism, and excretion
PDB: protein databank
RMSD: root-mean-square deviation
NAG: N-acetyl-D-glucosamine
NDG: 2-(acetylamino)-2-deoxy-A-D-glucopyranose
FUC: alpha-L-fucose
IFD: induced-fit docking

## LINKS TO ASSOCIATED CONTENT

Data associated with molecular dynamics simulations, TEST and DEREK NEXUS results, as well as fragments used to build the library and top ligands resulting from molecular docking, can be found on the following link: To be provided upon acceptance.

## Supporting Information

**Figure 1:**
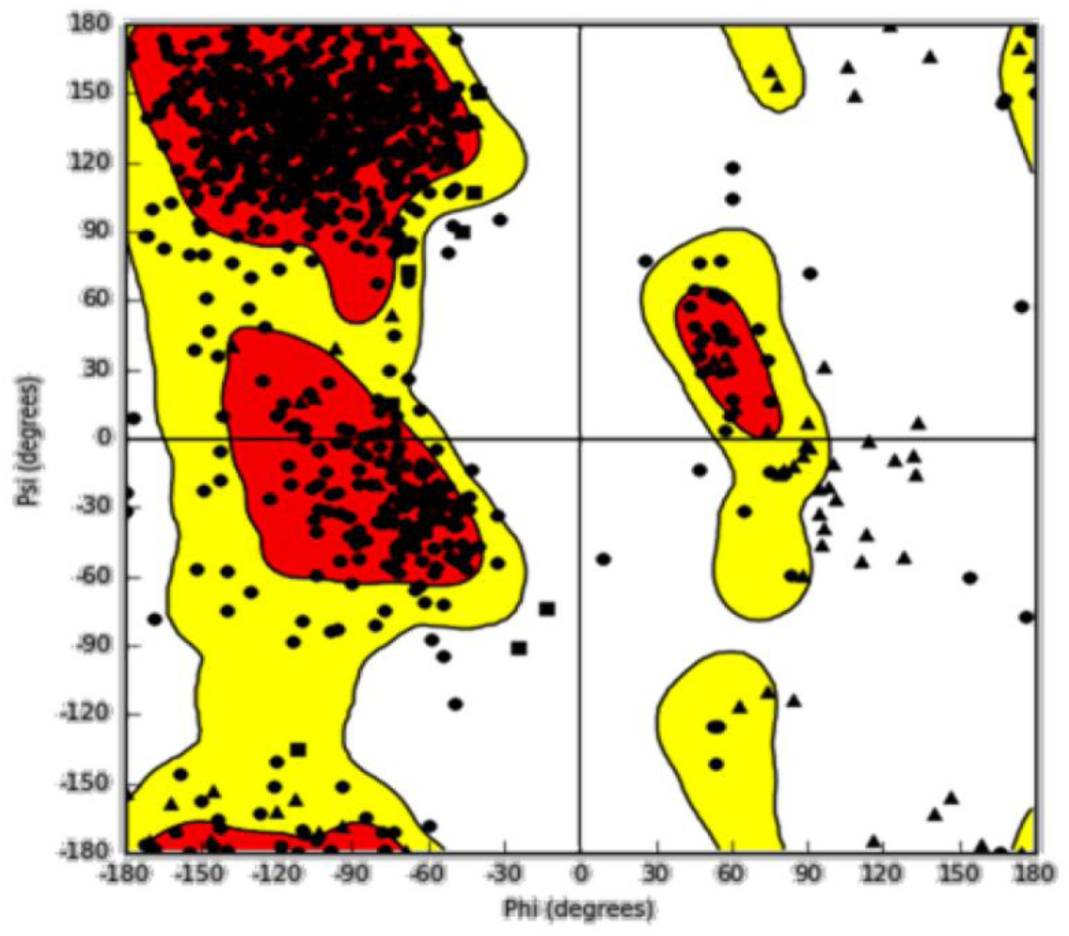
Ramachandran plot for 1GC1 pdb file preparation.

**Figure 2:**
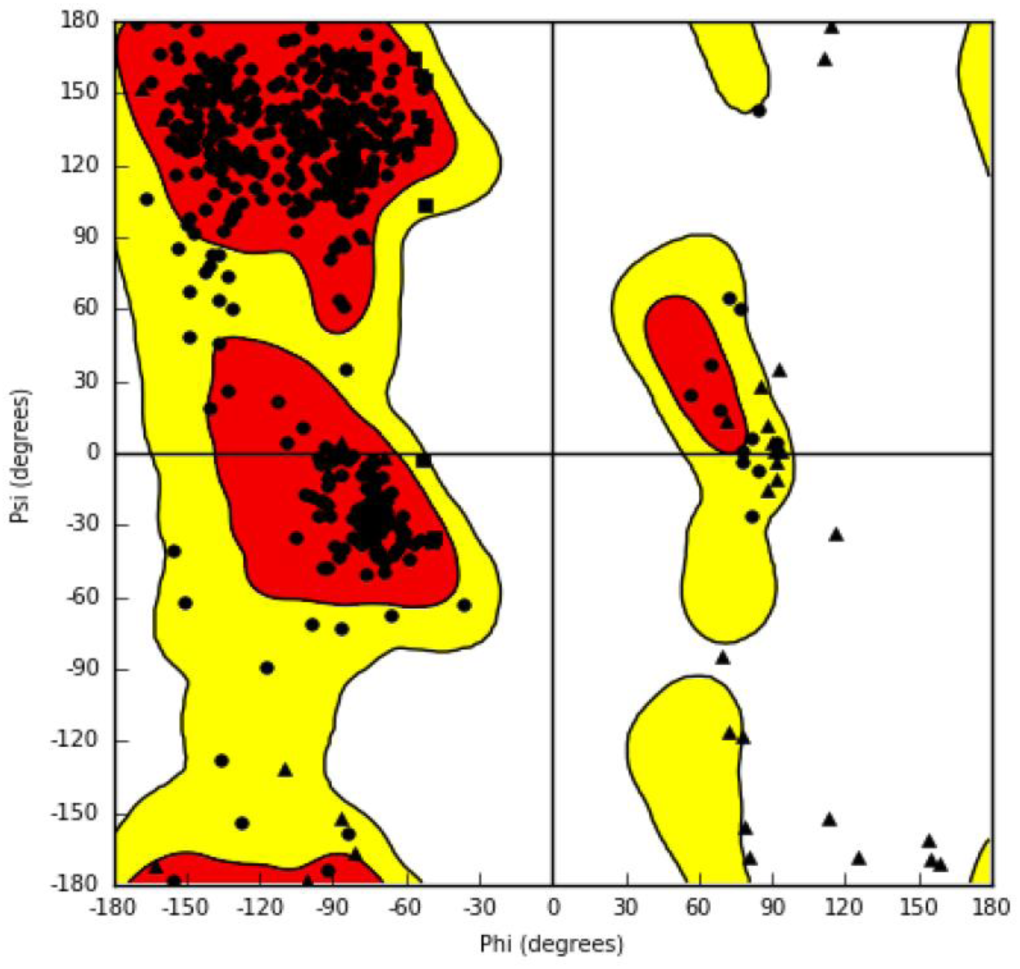
Ramachandran plot for 1G9M pdb file preparation.

**Table 1:** Consensus docking scores and first ranking values

| Ligand | DS | XP | DS | XP | LE<br>1gc1 | BC<br>1gc1 | LE | BC | Total | Rankin<br>g |
| --- | --- | --- | --- | --- | --- | --- | --- | --- | --- | --- |
|  | 1gc | 1gc | 1g9 | 1g9 |  |  | 1g9 | 1g9 |  |  |
|  | 1 | 1 | m | m |  |  | m | m |  |  |
| 17 | 8.64 | 8.64 | 8.27 | 8.27 | 10.45 | 10.41 | 10.57 | 10.57 | 37.90 | 1 |
| 18 | 8.64 | 8.64 | 8.27 | 8.27 | 10.44 | 10.40 | 10.55 | 10.55 | 37.87 | 2 |
| 6 | 8.92 | 8.92 | 8.49 | 8.49 | 10.03 | 10.03 | 10.43 | 10.43 | 37.87 | 3 |
| 94 | 8.11 | 8.11 | 8.04 | 8.04 | 10.77 | 10.77 | 10.85 | 10.85 | 37.77 | 4 |
| 95 | 8.11 | 8.11 | 8.04 | 8.04 | 10.78 | 10.78 | 10.83 | 10.83 | 37.76 | 5 |
| 96 | 8.11 | 8.11 | 8.04 | 8.04 | 10.73 | 10.73 | 10.86 | 10.86 | 37.74 | 6 |
| 22 | 8.60 | 8.60 | 7.62 | 7.62 | 10.04 | 10.04 | 10.36 | 10.36 | 36.62 | 7 |
| 121 | 8.03 | 8.03 | 7.41 | 7.41 | 10.45 | 10.45 | 10.43 | 10.43 | 36.31 | 8 |
| 64 | 8.24 | 8.48 | 7.44 | 7.44 | 10.16 | 10.16 | 10.35 | 10.35 | 36.31 | 9 |
| **65** | 8.24 | 8.48 | 7.44 | 7.44 | 10.16 | 10.16 | 10.35 | 10.35 | 36.31 | **9** |
| **122** | 8.03 | 8.03 | 7.41 | 7.41 | 10.45 | 10.45 | 10.41 | 10.41 | 36.29 | **11** |
| **32** | 8.42 | 8.42 | 7.62 | 7.62 | 9.67 | 9.67 | 10.02 | 10.02 | 35.73 | **12** |
| **60** | 8.24 | 8.24 | 6.49 | 6.49 | 10.26 | 10.26 | 10.64 | 10.64 | 35.63 | **13** |
| **160** | 7.92 | 7.92 | 6.46 | 6.46 | 10.25 | 10.25 | 10.71 | 10.71 | 35.33 | **14** |
| **88** | 8.14 | 8.49 | 8.61 | 8.61 | 9.09 | 9.09 | 9.52 | 9.08 | 35.31 | **15** |
| **267** | 7.71 | 7.85 | 6.93 | 6.93 | 10.03 | 10.03 | 10.37 | 10.37 | 35.11 | **16** |
| **63** | 8.24 | 8.24 | 7.76 | 7.76 | 9.70 | 9.70 | 9.37 | 9.37 | 35.07 | **17** |
| **238** | 7.75 | 7.75 | 6.76 | 6.76 | 10.27 | 10.27 | 10.28 | 10.28 | 35.06 | **18** |
| **67** | 8.22 | 8.22 | 7.77 | 7.77 | 9.71 | 9.71 | 9.35 | 9.35 | 35.06 | **19** |
| **59** | 8.24 | 8.83 | 8.02 | 8.02 | 8.74 | 8.74 | 9.66 | 9.66 | 34.96 | **20** |
| **56** | 8.26 | 8.39 | 7.90 | 7.90 | 9.60 | 9.60 | 9.09 | 9.09 | 34.91 | **21** |
| **255** | 7.73 | 7.73 | 6.73 | 6.73 | 9.97 | 9.97 | 10.44 | 10.44 | 34.87 | **22** |
| **253** | 7.74 | 8.01 | 7.41 | 7.41 | 10.09 | 10.09 | 9.47 | 9.47 | 34.84 | **23** |
| **105** | 8.09 | 8.43 | 8.02 | 8.02 | 9.28 | 9.28 | 9.26 | 9.26 | 34.82 | **24** |
| **106** | 8.09 | 8.43 | 8.02 | 8.02 | 9.28 | 9.28 | 9.26 | 9.26 | 34.82 | **24** |
| **129** | 8.00 | 8.03 | 7.46 | 7.46 | 9.45 | 9.45 | 9.81 | 9.81 | 34.74 | **26** |
| **195** | 7.85 | 7.85 | 7.71 | 7.71 | 9.53 | 9.53 | 9.61 | 9.61 | 34.70 | **27** |
| **225** | 7.78 | 7.78 | 4.80 | 4.80 | 10.86 | 10.86 | 11.23 | 11.23 | 34.67 | **28** |
| **52** | 8.28 | 8.61 | 7.57 | 7.57 | 9.48 | 9.48 | 9.15 | 9.15 | 34.64 | **29** |
| **51** | 8.28 | 8.61 | 7.57 | 7.57 | 9.47 | 9.47 | 9.15 | 9.15 | 34.63 | **30** |
| **184** | 7.88 | 7.88 | 5.89 | 5.89 | 10.51 | 10.51 | 10.35 | 10.35 | 34.63 | **31** |
| **194** | 7.85 | 7.85 | 7.71 | 7.71 | 9.53 | 9.53 | 9.51 | 9.51 | 34.60 | **32** |
| **117** | 8.05 | 8.05 | 6.98 | 6.98 | 9.86 | 9.86 | 9.70 | 9.70 | 34.59 | **33** |
| **118** | 8.05 | 8.05 | 6.98 | 6.98 | 9.86 | 9.86 | 9.70 | 9.70 | 34.59 | **33** |
| **48** | 8.30 | 8.32 | 4.50 | 4.50 | 10.77 | 10.77 | 10.99 | 10.99 | 34.56 | 35 |
| **85** | 8.14 | 8.14 | 6.83 | 6.83 | 9.62 | 9.62 | 9.97 | 9.97 | 34.56 | 36 |
| **153** | 7.94 | 8.27 | 7.66 | 7.66 | 9.43 | 9.30 | 9.51 | 9.36 | 34.56 | 37 |
| **73** | 8.17 | 8.23 | 6.95 | 6.95 | 9.73 | 9.73 | 9.65 | 9.65 | 34.52 | 38 |
| **74** | 8.17 | 8.23 | 6.95 | 6.95 | 9.73 | 9.73 | 9.65 | 9.65 | 34.52 | 38 |
| **82** | 8.16 | 8.30 | 7.59 | 7.59 | 9.41 | 9.27 | 9.30 | 9.30 | 34.46 | 40 |
| **66** | 8.23 | 8.23 | 8.04 | 8.04 | 8.93 | 8.72 | 9.33 | 9.33 | 34.43 | 41 |
| **261** | 7.72 | 7.72 | 7.09 | 7.09 | 10.27 | 10.27 | 9.47 | 9.21 | 34.42 | 42 |
| **78** | 8.16 | 8.20 | 6.71 | 6.71 | 9.94 | 9.94 | 9.59 | 9.59 | 34.42 | 43 |
| **190** | 7.86 | 7.86 | 7.98 | 7.98 | 9.15 | 9.15 | 9.39 | 9.20 | 34.29 | 44 |
| **39** | 8.37 | 8.37 | 7.60 | 7.60 | 9.23 | 9.21 | 9.10 | 9.10 | 34.29 | 45 |
| **191** | 7.86 | 7.86 | 7.82 | 7.82 | 9.46 | 9.46 | 9.19 | 9.06 | 34.27 | 46 |
| **9** | 8.83 | 8.84 | 6.22 | 6.22 | 9.46 | 9.46 | 9.71 | 9.71 | 34.22 | 47 |
| **155** | 7.93 | 8.21 | 8.02 | 8.02 | 9.12 | 9.12 | 9.01 | 9.01 | 34.22 | 48 |
| **202** | 7.83 | 7.83 | 8.24 | 8.24 | 9.04 | 9.04 | 9.06 | 9.06 | 34.17 | 49 |
| **209** | 7.81 | 7.81 | 6.44 | 6.44 | 9.81 | 9.81 | 10.09 | 10.09 | 34.15 | 50 |
| **125** | 8.02 | 8.02 | 7.73 | 7.73 | 8.96 | 8.96 | 9.40 | 9.40 | 34.11 | 51 |
| **54** | 8.27 | 8.27 | 7.30 | 7.30 | 9.14 | 9.14 | 9.39 | 9.39 | 34.09 | 52 |
| **2** | 8.99 | 9.01 | 6.19 | 6.19 | 9.17 | 8.99 | 9.78 | 9.78 | 34.05 | 53 |
| **234** | 7.76 | 8.40 | 6.41 | 6.41 | 9.61 | 9.61 | 9.87 | 9.87 | 33.98 | 54 |
| **157** | 7.93 | 8.34 | 7.11 | 7.11 | 9.23 | 9.23 | 9.50 | 9.50 | 33.97 | 55 |
| **45** | 8.32 | 8.32 | 7.78 | 7.78 | 8.83 | 8.83 | 9.04 | 9.04 | 33.97 | 56 |
| **46** | 8.32 | 8.32 | 7.78 | 7.78 | 8.83 | 8.83 | 9.04 | 9.04 | 33.97 | 56 |
| **236** | 7.76 | 7.76 | 3.27 | 3.27 | 11.46 | 11.46 | 11.47 | 11.47 | 33.96 | 58 |
| **196** | 7.85 | 7.87 | 7.49 | 7.49 | 9.47 | 9.47 | 9.10 | 9.10 | 33.92 | 59 |
| **178** | 7.89 | 7.89 | 5.71 | 5.71 | 10.25 | 10.25 | 10.04 | 10.04 | 33.89 | 60 |
| **179** | 7.89 | 7.89 | 5.71 | 5.71 | 10.25 | 10.25 | 10.04 | 10.04 | 33.89 | 60 |
| **19** | 8.63 | 8.63 | 6.38 | 6.38 | 9.12 | 9.12 | 9.70 | 9.70 | 33.82 | 62 |
| **24** | 8.54 | 8.54 | 7.24 | 7.24 | 9.11 | 9.11 | 8.89 | 8.89 | 33.78 | 63 |
| **31** | 8.46 | 8.46 | 6.77 | 6.77 | 9.06 | 8.85 | 9.55 | 9.55 | 33.74 | 64 |
| **87** | 8.14 | 8.14 | 5.85 | 5.85 | 9.93 | 9.93 | 9.81 | 9.81 | 33.73 | 65 |
| **68** | 8.20 | 8.53 | 6.73 | 6.73 | 9.23 | 9.00 | 9.52 | 9.52 | 33.73 | 66 |
| **108** | 8.08 | 8.08 | 7.76 | 7.76 | 8.98 | 8.82 | 9.33 | 8.57 | 33.69 | 67 |
| **30** | 8.46 | 8.46 | 6.03 | 6.03 | 9.62 | 9.62 | 9.46 | 9.46 | 33.57 | 68 |
| **103** | 8.09 | 8.09 | 3.92 | 3.92 | 10.70 | 10.70 | 10.84 | 10.84 | 33.56 | 69 |
| **104** | 8.09 | 8.09 | 3.92 | 3.92 | 10.70 | 10.70 | 10.84 | 10.84 | 33.56 | 69 |
| **127** | 8.01 | 8.39 | 7.10 | 7.10 | 9.27 | 9.27 | 8.97 | 8.97 | 33.54 | 71 |
| **115** | 8.07 | 8.07 | 6.38 | 6.38 | 9.47 | 9.47 | 9.62 | 9.62 | 33.54 | 72 |
| **131** | 7.99 | 8.27 | 3.67 | 3.67 | 10.61 | 10.61 | 11.07 | 11.07 | 33.48 | 73 |
| **35** | 8.40 | 8.40 | 8.10 | 8.10 | 8.43 | 8.43 | 8.49 | 8.49 | 33.42 | 74 |
| **213** | 7.81 | 7.84 | 7.26 | 7.26 | 9.26 | 9.26 | 9.07 | 9.07 | 33.42 | 75 |
| **36** | 8.38 | 8.42 | 5.51 | 5.51 | 9.61 | 9.61 | 9.85 | 9.85 | 33.37 | 76 |
| **12** | 8.77 | 8.77 | 6.89 | 6.89 | 8.62 | 8.62 | 9.01 | 9.01 | 33.29 | 77 |
| **44** | 8.32 | 8.32 | 7.01 | 7.01 | 8.93 | 8.93 | 9.15 | 8.87 | 33.27 | 78 |
| **40** | 8.36 | 8.36 | 6.21 | 6.21 | 9.02 | 9.02 | 9.64 | 9.64 | 33.22 | 79 |
| **75** | 8.17 | 8.17 | 6.60 | 6.60 | 8.98 | 8.98 | 9.44 | 9.44 | 33.19 | 80 |
| **76** | 8.17 | 8.17 | 6.60 | 6.60 | 8.98 | 8.98 | 9.44 | 9.44 | 33.19 | 80 |
| **49** | 8.29 | 8.29 | 3.35 | 3.35 | 10.53 | 10.53 | 10.99 | 10.99 | 33.16 | 82 |
| **47** | 8.30 | 8.31 | 6.34 | 6.34 | 9.31 | 9.31 | 9.17 | 9.17 | 33.12 | 83 |
| **140** | 7.97 | 7.97 | 6.46 | 6.46 | 9.40 | 9.40 | 9.27 | 9.27 | 33.11 | 84 |
| 141 | 7.97 | 7.97 | 6.46 | 6.46 | 9.40 | 9.40 | 9.27 | 9.27 | 33.11 | 84 |
| 112 | 8.08 | 8.08 | 7.65 | 7.65 | 8.58 | 8.58 | 8.85 | 8.74 | 33.10 | 86 |
| 230 | 7.77 | 7.79 | 5.96 | 5.96 | 9.81 | 9.81 | 9.54 | 9.54 | 33.09 | 87 |
| 97 | 8.11 | 8.11 | 6.70 | 6.70 | 9.01 | 9.01 | 9.26 | 9.26 | 33.08 | 88 |
| 55 | 8.26 | 8.29 | 6.28 | 6.28 | 9.24 | 9.24 | 9.27 | 9.27 | 33.07 | 89 |
| 177 | 7.89 | 7.92 | 6.27 | 6.27 | 9.15 | 9.15 | 9.72 | 9.72 | 33.05 | 90 |
| 81 | 8.16 | 8.16 | 8.00 | 8.00 | 8.55 | 8.55 | 8.33 | 8.33 | 33.03 | 91 |
| 259 | 7.72 | 7.72 | 7.82 | 7.82 | 8.39 | 8.39 | 9.07 | 9.07 | 33.00 | 92 |
| 86 | 8.14 | 8.14 | 4.43 | 4.43 | 10.51 | 10.51 | 9.98 | 9.79 | 32.97 | 93 |
| 113 | 8.08 | 8.08 | 7.30 | 7.30 | 8.69 | 8.69 | 8.90 | 8.90 | 32.97 | 94 |
| 42 | 8.34 | 8.34 | 7.99 | 7.99 | 8.38 | 8.38 | 8.25 | 8.25 | 32.96 | 95 |
| 53 | 8.27 | 8.27 | 5.22 | 5.22 | 9.66 | 9.66 | 9.77 | 9.77 | 32.91 | 96 |
| 20 | 8.61 | 8.83 | 3.79 | 3.79 | 10.05 | 10.05 | 10.34 | 10.34 | 32.90 | 97 |
| 246 | 7.75 | 8.27 | 7.30 | 7.30 | 8.74 | 8.69 | 8.88 | 8.88 | 32.90 | 98 |
| 139 | 7.98 | 8.25 | 3.38 | 3.38 | 10.75 | 10.75 | 10.52 | 10.52 | 32.76 | 99 |
| 21 | 8.60 | 8.82 | 3.64 | 3.64 | 10.05 | 10.05 | 10.34 | 10.34 | 32.74 | 100 |
| 181 | 7.89 | 7.99 | 6.93 | 6.93 | 9.09 | 8.94 | 8.85 | 8.85 | 32.73 | 101 |
| 219 | 7.79 | 7.79 | 3.44 | 3.44 | 10.63 | 10.63 | 10.86 | 10.86 | 32.72 | 102 |
| 152 | 7.94 | 7.98 | 6.08 | 6.08 | 9.25 | 9.25 | 9.42 | 9.42 | 32.71 | 103 |
| 92 | 8.12 | 8.12 | 7.90 | 7.90 | 8.32 | 8.32 | 8.37 | 8.37 | 32.71 | 104 |
| 93 | 8.12 | 8.12 | 7.90 | 7.90 | 8.32 | 8.32 | 8.37 | 8.37 | 32.71 | 104 |
| 13 | 8.72 | 8.72 | 6.45 | 6.45 | 8.54 | 8.54 | 8.99 | 8.99 | 32.70 | 106 |
| 14 | 8.72 | 8.72 | 6.45 | 6.45 | 8.54 | 8.54 | 8.99 | 8.99 | 32.70 | 106 |
| 26 | 8.48 | 8.48 | 5.98 | 5.98 | 9.01 | 8.98 | 9.24 | 9.24 | 32.69 | 108 |
| 50 | 8.28 | 8.28 | 6.54 | 6.54 | 8.80 | 8.80 | 9.03 | 9.03 | 32.65 | 109 |
| **187** | 7.88 | 7.88 | 6.56 | 6.56 | 8.79 | 8.65 | 9.49 | 9.49 | 32.64 | 110 |
| **244** | 7.75 | 7.76 | 5.98 | 5.98 | 9.43 | 9.43 | 9.43 | 9.43 | 32.59 | 111 |
| **245** | 7.75 | 7.76 | 5.98 | 5.98 | 9.43 | 9.43 | 9.43 | 9.43 | 32.59 | 111 |
| **185** | 7.88 | 7.88 | 7.50 | 7.50 | 8.38 | 8.38 | 8.81 | 8.81 | 32.57 | 113 |
| **239** | 7.75 | 7.75 | 6.18 | 6.18 | 9.35 | 9.35 | 9.26 | 9.26 | 32.54 | 114 |
| **98** | 8.11 | 8.11 | 3.42 | 3.42 | 10.38 | 10.38 | 10.63 | 10.63 | 32.54 | 115 |
| **99** | 8.11 | 8.11 | 3.42 | 3.42 | 10.38 | 10.38 | 10.63 | 10.63 | 32.54 | 115 |
| **188** | 7.87 | 8.01 | 7.34 | 7.34 | 8.55 | 8.55 | 8.82 | 8.51 | 32.49 | 117 |
| **166** | 7.91 | 7.91 | 2.66 | 2.66 | 10.88 | 10.88 | 11.04 | 11.04 | 32.49 | 118 |
| **58** | 8.25 | 8.25 | 7.10 | 7.10 | 8.44 | 8.44 | 8.68 | 8.68 | 32.47 | 119 |
| **258** | 7.73 | 7.77 | 5.23 | 5.23 | 9.54 | 9.54 | 9.94 | 9.94 | 32.45 | 120 |
| **130** | 7.99 | 7.99 | 4.42 | 4.42 | 10.25 | 10.25 | 9.95 | 9.63 | 32.45 | 121 |
| **212** | 7.81 | 7.81 | 7.21 | 7.21 | 8.62 | 8.62 | 8.86 | 8.76 | 32.45 | 122 |
| **124** | 8.02 | 8.11 | 6.40 | 6.40 | 8.92 | 8.92 | 9.05 | 9.05 | 32.44 | 123 |
| **25** | 8.50 | 8.50 | 7.58 | 7.58 | 7.95 | 7.95 | 8.40 | 8.40 | 32.43 | 124 |
| **154** | 7.93 | 7.93 | 4.89 | 4.89 | 9.94 | 9.94 | 9.65 | 9.65 | 32.41 | 125 |
| **260** | 7.72 | 7.72 | 3.23 | 3.23 | 10.59 | 10.59 | 10.82 | 10.82 | 32.36 | 126 |
| **41** | 8.34 | 8.34 | 6.95 | 6.95 | 8.14 | 8.14 | 8.92 | 8.92 | 32.35 | 127 |
| **111** | 8.08 | 8.14 | 8.12 | 8.12 | 7.63 | 7.63 | 8.46 | 8.46 | 32.32 | 128 |
| **176** | 7.89 | 8.48 | 3.67 | 3.67 | 10.06 | 9.91 | 10.43 | 10.42 | 32.27 | 129 |
| **143** | 7.96 | 7.97 | 5.98 | 5.98 | 8.93 | 8.93 | 9.37 | 9.37 | 32.24 | 130 |
| **144** | 7.96 | 7.97 | 5.98 | 5.98 | 8.93 | 8.93 | 9.37 | 9.37 | 32.24 | 130 |
| **110** | 8.08 | 8.26 | 7.85 | 7.85 | 8.05 | 8.05 | 8.17 | 8.17 | 32.24 | 132 |
| **83** | 8.15 | 8.16 | 5.10 | 5.10 | 9.73 | 9.73 | 9.24 | 9.24 | 32.23 | 133 |
| **84** | 8.15 | 8.16 | 5.10 | 5.10 | 9.73 | 9.73 | 9.24 | 9.24 | 32.23 | 133 |
| **214** | 7.80 | 7.80 | 8.45 | 8.45 | 8.00 | 8.00 | 7.97 | 7.94 | 32.21 | 135 |
| **182** | 7.89 | 7.89 | 4.60 | 4.60 | 9.74 | 9.74 | 9.97 | 9.97 | 32.20 | 136 |
| **70** | 8.20 | 8.20 | 6.14 | 6.14 | 8.60 | 8.60 | 9.25 | 9.25 | 32.19 | 137 |
| **268** | 7.71 | 7.71 | 6.37 | 6.37 | 9.28 | 9.24 | 8.83 | 8.83 | 32.17 | 138 |
| **43** | 8.33 | 8.33 | 5.98 | 5.98 | 8.93 | 8.93 | 8.93 | 8.93 | 32.16 | 139 |
| **138** | 7.98 | 8.63 | 3.47 | 3.47 | 10.15 | 10.15 | 10.17 | 10.17 | 32.10 | 140 |
| **189** | 7.87 | 7.87 | 6.61 | 6.61 | 8.44 | 8.44 | 9.16 | 9.16 | 32.07 | 141 |
| **241** | 7.75 | 7.75 | 7.06 | 7.06 | 8.44 | 8.44 | 8.80 | 8.80 | 32.05 | 142 |
| **242** | 7.75 | 7.75 | 7.06 | 7.06 | 8.44 | 8.44 | 8.80 | 8.80 | 32.05 | 142 |
| **180** | 7.89 | 7.89 | 7.54 | 7.54 | 8.21 | 8.21 | 8.40 | 8.40 | 32.04 | 144 |
| **226** | 7.78 | 7.78 | 4.59 | 4.59 | 9.69 | 9.69 | 9.97 | 9.97 | 32.03 | 145 |
| **34** | 8.41 | 8.41 | 5.74 | 5.74 | 8.78 | 8.78 | 9.06 | 9.06 | 31.99 | 146 |
| **33** | 8.41 | 8.41 | 5.74 | 5.74 | 8.76 | 8.76 | 9.06 | 9.06 | 31.97 | 147 |
| **7** | 8.88 | 8.88 | 4.40 | 4.40 | 9.44 | 9.44 | 9.50 | 8.96 | 31.96 | 148 |
| **132** | 7.99 | 8.07 | 5.42 | 5.42 | 8.85 | 8.85 | 9.63 | 9.63 | 31.93 | 149 |
| **198** | 7.84 | 7.84 | 7.20 | 7.20 | 8.30 | 8.30 | 8.58 | 8.58 | 31.92 | 150 |
| **172** | 7.90 | 7.90 | 5.07 | 5.07 | 9.40 | 9.40 | 9.52 | 9.52 | 31.89 | 151 |
| **8** | 8.87 | 8.92 | 5.53 | 5.53 | 8.67 | 8.36 | 8.93 | 8.93 | 31.86 | 152 |
| **217** | 7.80 | 7.80 | 4.00 | 4.00 | 10.20 | 10.03 | 9.94 | 9.94 | 31.85 | 153 |
| **107** | 8.09 | 8.21 | 6.47 | 6.47 | 8.41 | 8.41 | 8.79 | 8.79 | 31.82 | 154 |
| **146** | 7.96 | 7.96 | 7.92 | 7.92 | 7.83 | 7.83 | 8.11 | 8.11 | 31.82 | 155 |
| **4** | 8.98 | 8.98 | 4.45 | 4.45 | 9.18 | 9.18 | 9.34 | 8.95 | 31.75 | 156 |
| **199** | 7.84 | 7.84 | 6.94 | 6.94 | 8.48 | 8.48 | 8.70 | 8.28 | 31.75 | 157 |
| **3** | 8.98 | 8.98 | 4.45 | 4.45 | 9.17 | 9.17 | 9.35 | 8.94 | 31.74 | 158 |
| **120** | 8.04 | 8.06 | 4.88 | 4.88 | 9.23 | 9.23 | 9.69 | 9.45 | 31.73 | 159 |
| **23** | 8.59 | 9.15 | 4.95 | 4.95 | 8.72 | 8.72 | 9.19 | 9.19 | 31.72 | 160 |
| **77** | 8.17 | 8.17 | 7.22 | 7.22 | 7.84 | 7.84 | 8.49 | 8.49 | 31.72 | 161 |
| **273** | 7.70 | 7.70 | 3.70 | 3.70 | 10.32 | 10.32 | 9.99 | 9.99 | 31.71 | 162 |
| **247** | 7.75 | 7.75 | 7.13 | 7.13 | 7.98 | 7.94 | 8.87 | 8.87 | 31.70 | 163 |
| **197** | 7.85 | 7.85 | 7.35 | 7.35 | 7.98 | 7.98 | 8.50 | 8.50 | 31.68 | 164 |
| **193** | 7.85 | 7.85 | 5.41 | 5.41 | 9.64 | 9.28 | 9.01 | 8.90 | 31.68 | 165 |
| **252** | 7.74 | 7.78 | 4.46 | 4.46 | 9.49 | 9.49 | 9.90 | 9.90 | 31.61 | 166 |
| **37** | 8.38 | 8.38 | 6.45 | 6.45 | 8.13 | 8.13 | 8.67 | 8.62 | 31.60 | 167 |
| **142** | 7.97 | 7.97 | 3.82 | 3.82 | 9.45 | 9.45 | 10.32 | 10.32 | 31.57 | 168 |
| **192** | 7.85 | 7.85 | 6.20 | 6.20 | 8.70 | 8.70 | 8.76 | 8.76 | 31.51 | 169 |
| **69** | 8.20 | 8.27 | 5.60 | 5.60 | 8.87 | 8.82 | 8.80 | 8.80 | 31.48 | 170 |
| **149** | 7.96 | 7.96 | 6.56 | 6.56 | 8.45 | 8.45 | 8.45 | 8.45 | 31.42 | 171 |
| **224** | 7.78 | 8.06 | 4.18 | 4.18 | 9.61 | 9.61 | 9.69 | 9.69 | 31.40 | 172 |
| **147** | 7.96 | 7.98 | 6.15 | 6.15 | 8.41 | 8.41 | 8.81 | 8.81 | 31.34 | 173 |
| **271** | 7.71 | 8.03 | 3.66 | 3.66 | 10.00 | 10.00 | 9.80 | 9.80 | 31.33 | 174 |
| **148** | 7.96 | 7.98 | 6.15 | 6.15 | 8.41 | 8.41 | 8.80 | 8.80 | 31.33 | 175 |
| **133** | 7.98 | 8.01 | 6.47 | 6.47 | 8.34 | 8.34 | 8.38 | 8.38 | 31.18 | 176 |
| **134** | 7.98 | 8.01 | 6.47 | 6.47 | 8.34 | 8.34 | 8.38 | 8.38 | 31.18 | 176 |
| **135** | 7.98 | 7.98 | 6.17 | 6.17 | 8.12 | 8.02 | 8.96 | 8.96 | 31.18 | 178 |
| **265** | 7.71 | 7.71 | 6.41 | 6.41 | 8.09 | 8.00 | 9.01 | 9.01 | 31.17 | 179 |
| **205** | 7.82 | 7.82 | 6.29 | 6.29 | 8.18 | 8.18 | 8.82 | 8.82 | 31.11 | 180 |
| **208** | 7.81 | 7.81 | 8.09 | 8.09 | 7.63 | 7.59 | 7.64 | 7.51 | 31.09 | 181 |
| **229** | 7.77 | 7.89 | 5.05 | 5.05 | 9.10 | 9.10 | 9.11 | 9.04 | 31.06 | 182 |
| **250** | 7.74 | 7.74 | 7.19 | 7.19 | 7.78 | 7.78 | 8.35 | 8.35 | 31.06 | 183 |
| **270** | 7.71 | 7.71 | 6.10 | 6.10 | 8.40 | 8.40 | 8.82 | 8.82 | 31.03 | 184 |
| 158 | 7.92 | 7.92 | 3.33 | 3.33 | 9.76 | 9.76 | 10.00 | 10.00 | 31.01 | 185 |
| 123 | 8.03 | 8.03 | 2.57 | 2.57 | 9.82 | 9.82 | 10.54 | 10.52 | 30.94 | 186 |
| 248 | 7.74 | 7.74 | 5.70 | 5.70 | 8.77 | 8.53 | 8.91 | 8.69 | 30.89 | 187 |
| 249 | 7.74 | 7.74 | 5.70 | 5.70 | 8.77 | 8.53 | 8.91 | 8.69 | 30.89 | 187 |
| 79 | 8.16 | 8.16 | 6.55 | 6.55 | 7.81 | 7.81 | 8.35 | 8.35 | 30.87 | 189 |
| 223 | 7.78 | 8.29 | 3.18 | 3.18 | 9.84 | 9.84 | 9.73 | 9.73 | 30.78 | 190 |
| 210 | 7.81 | 7.81 | 2.65 | 2.65 | 10.08 | 10.08 | 10.25 | 10.23 | 30.78 | 191 |
| 231 | 7.77 | 8.10 | 3.36 | 3.36 | 9.68 | 9.68 | 9.80 | 9.80 | 30.77 | 192 |
| 71 | 8.20 | 8.20 | 7.58 | 7.58 | 7.33 | 7.33 | 7.66 | 7.66 | 30.77 | 193 |
| 114 | 8.07 | 8.14 | 7.06 | 7.06 | 7.57 | 7.57 | 8.03 | 8.03 | 30.76 | 194 |
| 161 | 7.91 | 7.91 | 3.89 | 3.89 | 9.32 | 9.32 | 9.63 | 9.63 | 30.76 | 195 |
| 28 | 8.48 | 9.04 | 4.93 | 4.93 | 8.07 | 8.04 | 9.01 | 9.01 | 30.75 | 196 |
| 29 | 8.48 | 9.04 | 4.93 | 4.93 | 8.07 | 8.04 | 9.00 | 9.00 | 30.74 | 197 |
| 186 | 7.88 | 7.88 | 7.09 | 7.09 | 7.62 | 7.62 | 8.07 | 8.07 | 30.65 | 198 |
| 167 | 7.91 | 7.91 | 6.43 | 6.43 | 8.01 | 8.01 | 8.11 | 8.11 | 30.46 | 199 |
| 204 | 7.83 | 7.83 | 7.27 | 7.27 | 7.49 | 7.49 | 7.86 | 7.86 | 30.45 | 200 |
| 235 | 7.76 | 7.79 | 5.83 | 5.83 | 8.51 | 8.35 | 8.38 | 8.38 | 30.41 | 201 |
| 128 | 8.00 | 8.03 | 4.75 | 4.75 | 8.67 | 8.67 | 8.95 | 8.95 | 30.39 | 202 |
| 100 | 8.11 | 8.11 | 6.70 | 6.70 | 7.68 | 7.68 | 7.90 | 7.90 | 30.39 | 203 |
| 15 | 8.66 | 8.66 | 7.13 | 7.13 | 7.02 | 7.02 | 7.61 | 7.53 | 30.39 | 204 |
| 269 | 7.71 | 7.75 | 4.97 | 4.97 | 8.68 | 8.68 | 9.00 | 9.00 | 30.38 | 205 |
| 207 | 7.82 | 7.82 | 4.83 | 4.83 | 8.76 | 8.76 | 8.91 | 8.91 | 30.31 | 206 |
| 164 | 7.91 | 8.48 | 4.96 | 4.96 | 8.81 | 8.81 | 8.33 | 8.33 | 30.30 | 207 |
| 233 | 7.77 | 7.77 | 6.27 | 6.27 | 7.77 | 7.77 | 8.45 | 8.45 | 30.26 | 208 |
| 165 | 7.91 | 8.48 | 4.96 | 4.96 | 8.80 | 8.80 | 8.34 | 8.24 | 30.25 | 209 |
| **119** | 8.05 | 8.31 | 7.34 | 7.34 | 7.29 | 7.29 | 7.41 | 7.41 | 30.22 | 210 |
| **151** | 7.95 | 7.95 | 6.32 | 6.32 | 7.76 | 7.76 | 8.19 | 8.19 | 30.22 | 210 |
| **137** | 7.98 | 7.98 | 7.22 | 7.22 | 7.39 | 7.39 | 7.62 | 7.62 | 30.21 | 212 |
| **150** | 7.95 | 7.95 | 6.32 | 6.32 | 7.76 | 7.76 | 8.18 | 8.18 | 30.21 | 213 |
| **215** | 7.80 | 8.13 | 2.73 | 2.73 | 9.68 | 9.68 | 9.79 | 9.79 | 30.17 | 214 |
| **216** | 7.80 | 7.80 | 6.05 | 6.05 | 8.03 | 8.03 | 8.31 | 8.24 | 30.15 | 215 |
| **109** | 8.08 | 8.09 | 6.01 | 6.01 | 8.15 | 8.15 | 7.93 | 7.87 | 30.14 | 216 |
| **251** | 7.74 | 7.78 | 2.88 | 2.88 | 9.95 | 9.95 | 9.53 | 9.53 | 30.12 | 217 |
| **102** | 8.10 | 8.10 | 5.95 | 5.95 | 8.17 | 8.17 | 7.89 | 7.89 | 30.11 | 218 |
| **201** | 7.84 | 7.84 | 3.34 | 3.34 | 9.22 | 9.22 | 9.71 | 9.71 | 30.11 | 219 |
| **228** | 7.77 | 7.77 | 6.03 | 6.03 | 7.91 | 7.91 | 8.35 | 8.35 | 30.07 | 220 |
| **5** | 8.94 | 8.97 | 4.23 | 4.23 | 8.27 | 8.25 | 8.61 | 8.61 | 30.06 | 221 |
| **136** | 7.98 | 7.98 | 3.37 | 3.37 | 9.27 | 9.27 | 9.42 | 9.42 | 30.03 | 222 |
| **169** | 7.90 | 7.90 | 2.04 | 2.04 | 10.04 | 10.04 | 9.99 | 9.99 | 29.97 | 223 |
| **168** | 7.90 | 7.90 | 2.04 | 2.04 | 10.04 | 10.04 | 9.96 | 9.96 | 29.94 | 224 |
| **61** | 8.24 | 8.24 | 2.03 | 2.03 | 9.65 | 9.65 | 9.99 | 9.99 | 29.91 | 225 |
| **62** | 8.24 | 8.24 | 2.03 | 2.03 | 9.65 | 9.65 | 9.99 | 9.99 | 29.91 | 225 |
| **173** | 7.89 | 7.89 | 6.87 | 6.87 | 7.45 | 7.45 | 7.67 | 7.67 | 29.89 | 227 |
| **174** | 7.89 | 7.89 | 6.87 | 6.87 | 7.45 | 7.45 | 7.67 | 7.67 | 29.89 | 227 |
| **175** | 7.89 | 7.89 | 6.87 | 6.87 | 7.45 | 7.45 | 7.67 | 7.67 | 29.89 | 227 |
| **227** | 7.78 | 7.88 | 4.67 | 4.67 | 8.59 | 8.59 | 8.76 | 8.76 | 29.84 | 230 |
| **91** | 8.13 | 8.13 | 2.10 | 2.10 | 9.58 | 9.58 | 9.91 | 9.91 | 29.71 | 231 |
| **274** | 7.70 | 8.34 | 0.00 | 0.00 | 10.74 | 10.74 | 10.93 | 10.93 | 29.69 | 232 |
| **72** | 8.18 | 8.18 | 1.91 | 1.91 | 9.60 | 9.60 | 10.11 | 9.82 | 29.66 | 233 |
| **220** | 7.79 | 7.79 | 0.76 | 0.76 | 10.56 | 10.56 | 10.46 | 10.46 | 29.57 | 234 |
| **183** | 7.89 | 7.89 | 2.48 | 2.48 | 9.55 | 9.55 | 9.65 | 9.65 | 29.57 | 235 |
| **237** | 7.76 | 7.76 | 7.13 | 7.13 | 7.33 | 7.33 | 7.32 | 7.32 | 29.54 | 236 |
| **101** | 8.10 | 8.43 | 3.18 | 3.18 | 9.15 | 9.15 | 8.88 | 8.88 | 29.47 | 237 |
| **203** | 7.83 | 8.33 | 1.56 | 1.56 | 9.95 | 9.50 | 10.11 | 10.11 | 29.47 | 238 |
| **254** | 7.73 | 7.73 | 4.89 | 4.89 | 8.19 | 8.19 | 8.60 | 8.60 | 29.41 | 239 |
| **159** | 7.92 | 7.92 | 2.99 | 2.99 | 9.46 | 9.46 | 9.01 | 8.99 | 29.37 | 240 |
| **116** | 8.06 | 8.08 | 4.76 | 4.76 | 8.26 | 8.26 | 8.27 | 8.23 | 29.34 | 241 |
| **263** | 7.71 | 7.71 | 4.59 | 4.59 | 8.48 | 8.48 | 8.56 | 8.56 | 29.34 | 242 |
| **264** | 7.71 | 7.71 | 4.57 | 4.57 | 8.48 | 8.48 | 8.56 | 8.56 | 29.32 | 243 |
| **262** | 7.71 | 7.71 | 2.65 | 2.65 | 9.46 | 9.36 | 9.47 | 9.47 | 29.24 | 244 |
| **232** | 7.77 | 7.77 | 5.76 | 5.76 | 7.77 | 7.77 | 7.93 | 7.93 | 29.23 | 245 |
| **218** | 7.79 | 7.79 | 1.31 | 1.31 | 9.87 | 9.87 | 10.13 | 10.13 | 29.10 | 246 |
| **206** | 7.82 | 7.82 | 5.49 | 5.49 | 7.80 | 7.80 | 7.93 | 7.92 | 29.03 | 247 |
| **90** | 8.13 | 8.13 | 0.88 | 0.88 | 9.82 | 9.75 | 10.25 | 9.93 | 28.89 | 248 |
| **38** | 8.37 | 8.37 | 1.89 | 1.89 | 9.26 | 9.26 | 9.34 | 9.34 | 28.86 | 249 |
| **211** | 7.81 | 7.81 | 2.82 | 2.82 | 8.82 | 8.82 | 9.06 | 9.06 | 28.51 | 250 |
| **145** | 7.96 | 7.96 | 3.28 | 3.28 | 8.41 | 8.41 | 8.42 | 8.42 | 28.07 | 251 |
| **200** | 7.84 | 7.92 | 2.78 | 2.78 | 8.67 | 8.67 | 8.74 | 8.33 | 27.86 | 252 |
| **10** | 8.82 | 8.82 | 0.00 | 0.00 | 9.24 | 9.02 | 9.82 | 9.82 | 27.77 | 253 |
| **11** | 8.82 | 8.82 | 0.00 | 0.00 | 9.07 | 9.02 | 9.82 | 9.82 | 27.69 | 254 |
| **1** | 9.09 | 9.09 | 0.00 | 0.00 | 8.98 | 8.98 | 9.58 | 9.58 | 27.65 | 255 |
| **126** | 8.02 | 8.02 | 1.74 | 1.74 | 8.80 | 8.80 | 9.20 | 8.94 | 27.62 | 256 |
| **57** | 8.26 | 8.41 | 0.00 | 0.00 | 9.42 | 9.42 | 9.68 | 9.68 | 27.43 | 257 |
| **266** | 7.71 | 7.78 | 1.21 | 1.21 | 9.13 | 9.13 | 9.13 | 9.13 | 27.21 | 258 |
| **163** | 7.91 | 8.46 | 0.00 | 0.00 | 9.28 | 9.28 | 9.57 | 9.57 | 27.04 | 259 |
| **162** | 7.91 | 8.46 | 0.00 | 0.00 | 9.27 | 9.27 | 9.57 | 9.57 | 27.03 | 260 |
| **243** | 7.75 | 7.75 | 2.63 | 2.63 | 8.03 | 8.03 | 8.51 | 8.51 | 26.92 | 261 |
| **170** | 7.90 | 7.90 | 0.31 | 0.31 | 9.16 | 9.16 | 9.36 | 9.36 | 26.73 | 262 |
| **171** | 7.90 | 7.90 | 0.31 | 0.31 | 9.16 | 9.16 | 9.36 | 9.36 | 26.73 | 262 |
| **156** | 7.93 | 8.49 | 1.36 | 1.36 | 8.50 | 8.50 | 8.59 | 8.59 | 26.66 | 264 |
| **80** | 8.16 | 8.18 | 0.00 | 0.00 | 9.00 | 9.00 | 9.26 | 9.26 | 26.43 | 265 |
| **256** | 7.73 | 7.73 | 0.00 | 0.00 | 9.21 | 9.21 | 9.24 | 9.24 | 26.18 | 266 |
| **27** | 8.48 | 8.48 | 0.00 | 0.00 | 8.66 | 8.66 | 8.76 | 8.76 | 25.90 | 267 |
| **221** | 7.78 | 7.80 | 1.78 | 1.78 | 8.11 | 8.11 | 8.01 | 8.01 | 25.69 | 268 |
| **222** | 7.78 | 7.80 | 1.70 | 1.70 | 8.11 | 8.11 | 8.01 | 8.01 | 25.61 | 269 |
| **272** | 7.70 | 7.70 | 1.60 | 1.60 | 7.71 | 7.71 | 8.53 | 8.53 | 25.54 | 270 |
| **16** | 8.65 | 8.65 | 0.00 | 0.00 | 8.66 | 8.66 | 8.37 | 8.02 | 25.51 | 271 |
| **240** | 7.75 | 7.75 | 0.00 | 0.00 | 8.69 | 8.69 | 8.84 | 8.84 | 25.28 | 272 |
| **275** | 7.70 | 7.70 | 0.00 | 0.00 | 8.28 | 8.28 | 8.51 | 8.51 | 24.49 | 273 |
| **257** | 7.73 | 7.73 | 0.00 | 0.00 | 8.38 | 7.90 | 8.62 | 8.62 | 24.49 | 274 |
| **89** | 8.14 | 8.14 | 0.00 | 0.00 | 7.63 | 7.63 | 7.94 | 7.93 | 23.70 | 275 |

**Table 2:** Top 20 compounds IFD values and ranking.

| Ligand | IFD<br>ABS | Ranking |
| --- | --- | --- |
| **255** | 646.08 | 1 |
| **141** | 644.33 | 2 |
| **2** | 644.28 | 3 |
| **81** | 644.11 | 4 |
| **9** | 643.97 | 5 |
| **22** | 643.67 | 6 |
| **59** | 643.64 | 7 |
| **177** | 643.58 | 8 |
| **85** | 643.31 | 9 |
| **44** | 643.00 | 10 |
| **157** | 642.99 | 11 |
| **140** | 642.94 | 12 |
| **86** | 642.82 | 13 |
| **234** | 642.61 | 14 |
| **48** | 642.59 | 15 |
| **129** | 642.35 | 16 |
| **160** | 642.24 | 17 |
| **73** | 642.08 | 18 |
| **74** | 641.92 | 19 |
| **155** | 641.86 | 20 |

**Figure 3:**
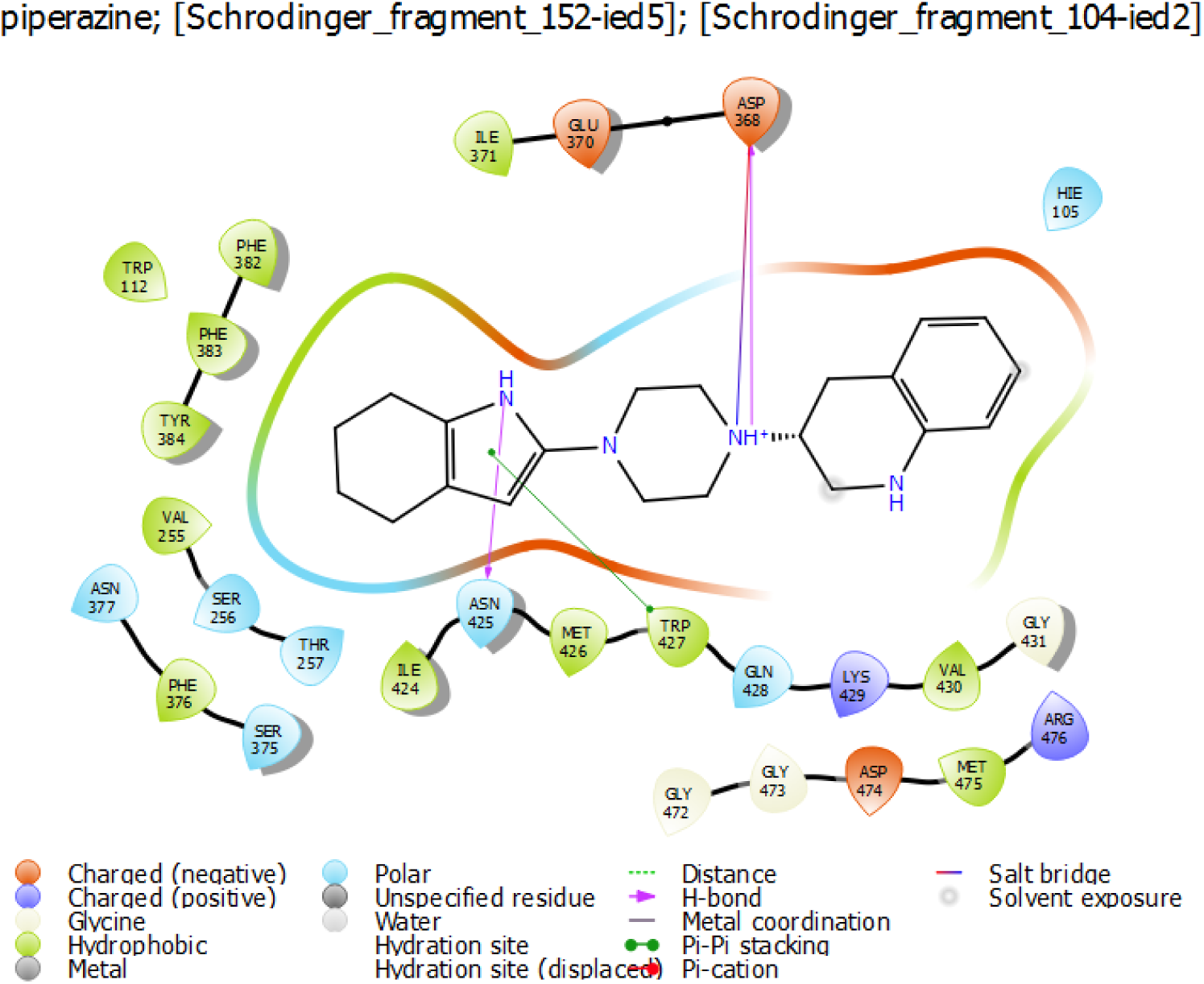
IFD interaction diagram for ligand 141.

**Figure 4:**
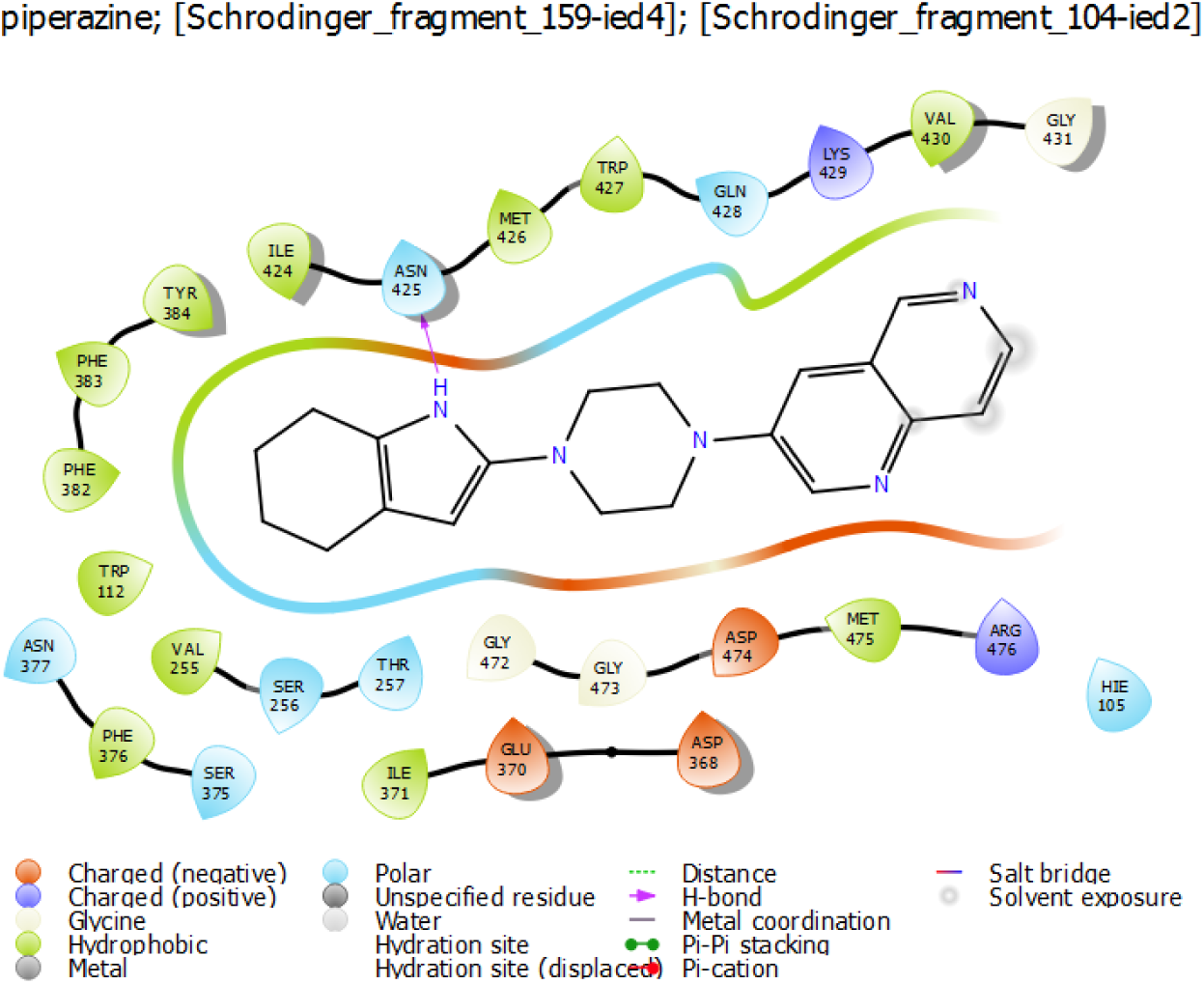
IFD interaction diagram for ligand 002.

**Figure 5:**
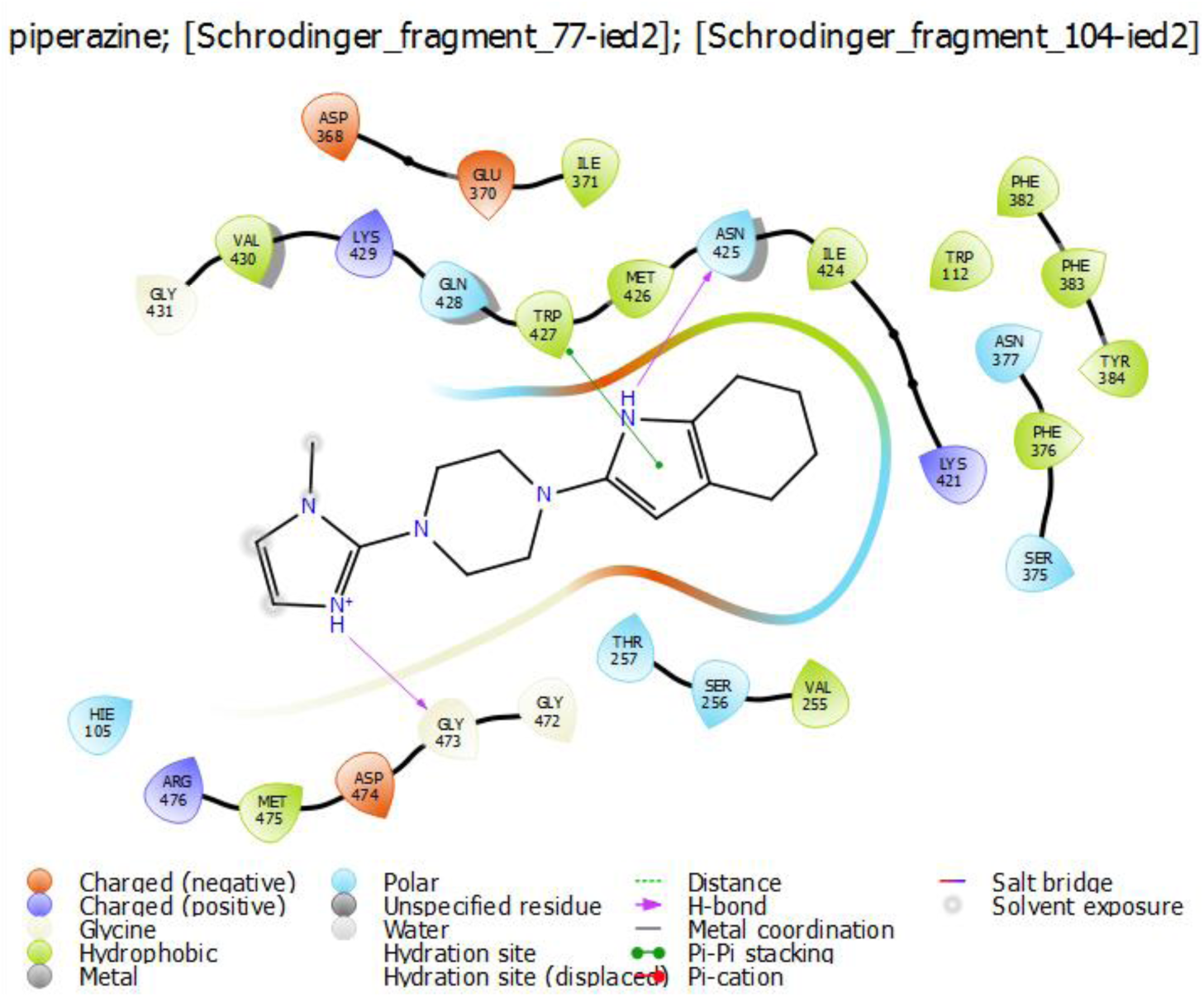
IFD interaction diagram for ligand 081.

**Figure 6:**
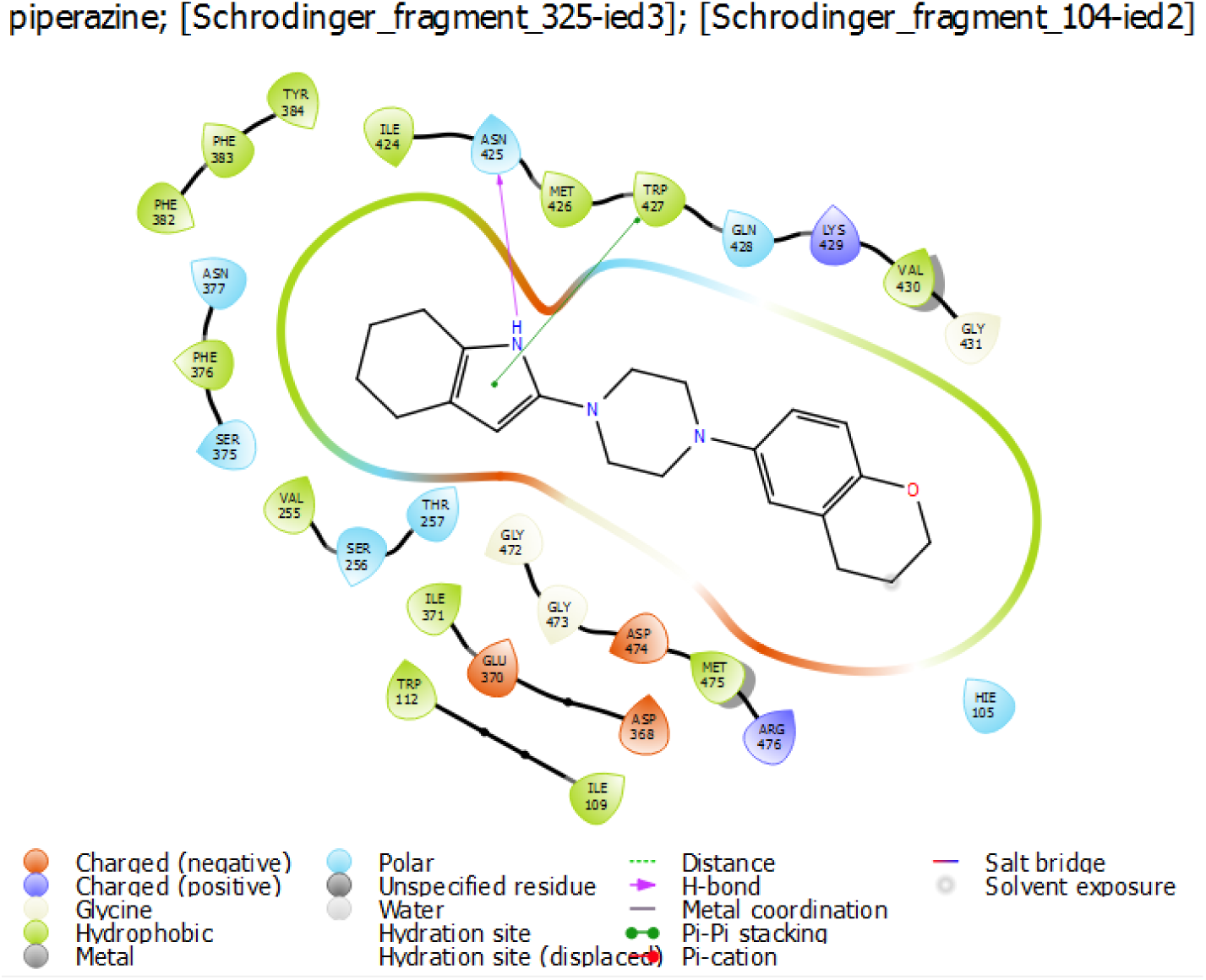
IFD interaction diagram for ligand 009.

**Figure 7:**
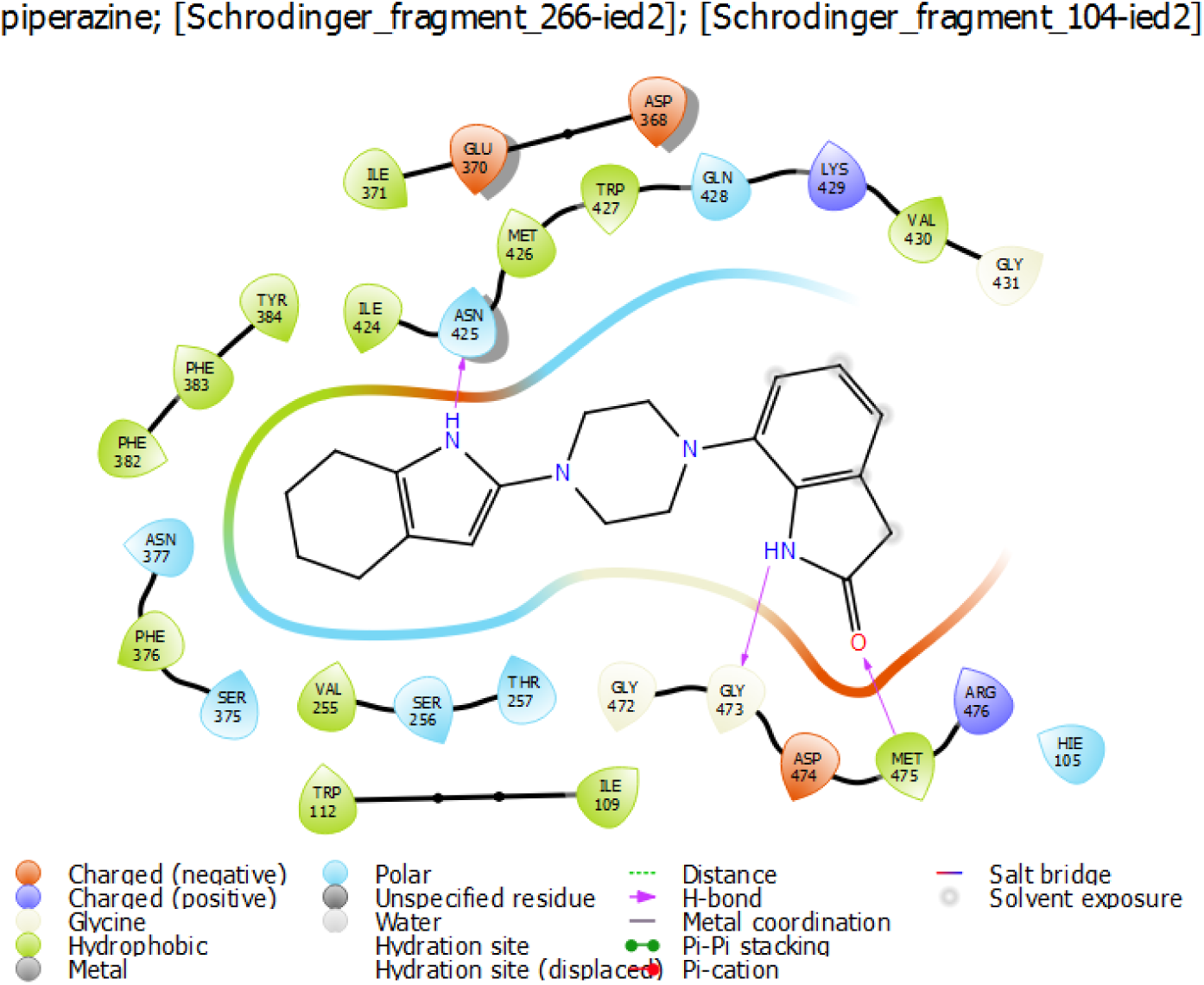
IFD interaction diagram for ligand 022.

**Figure 8:**
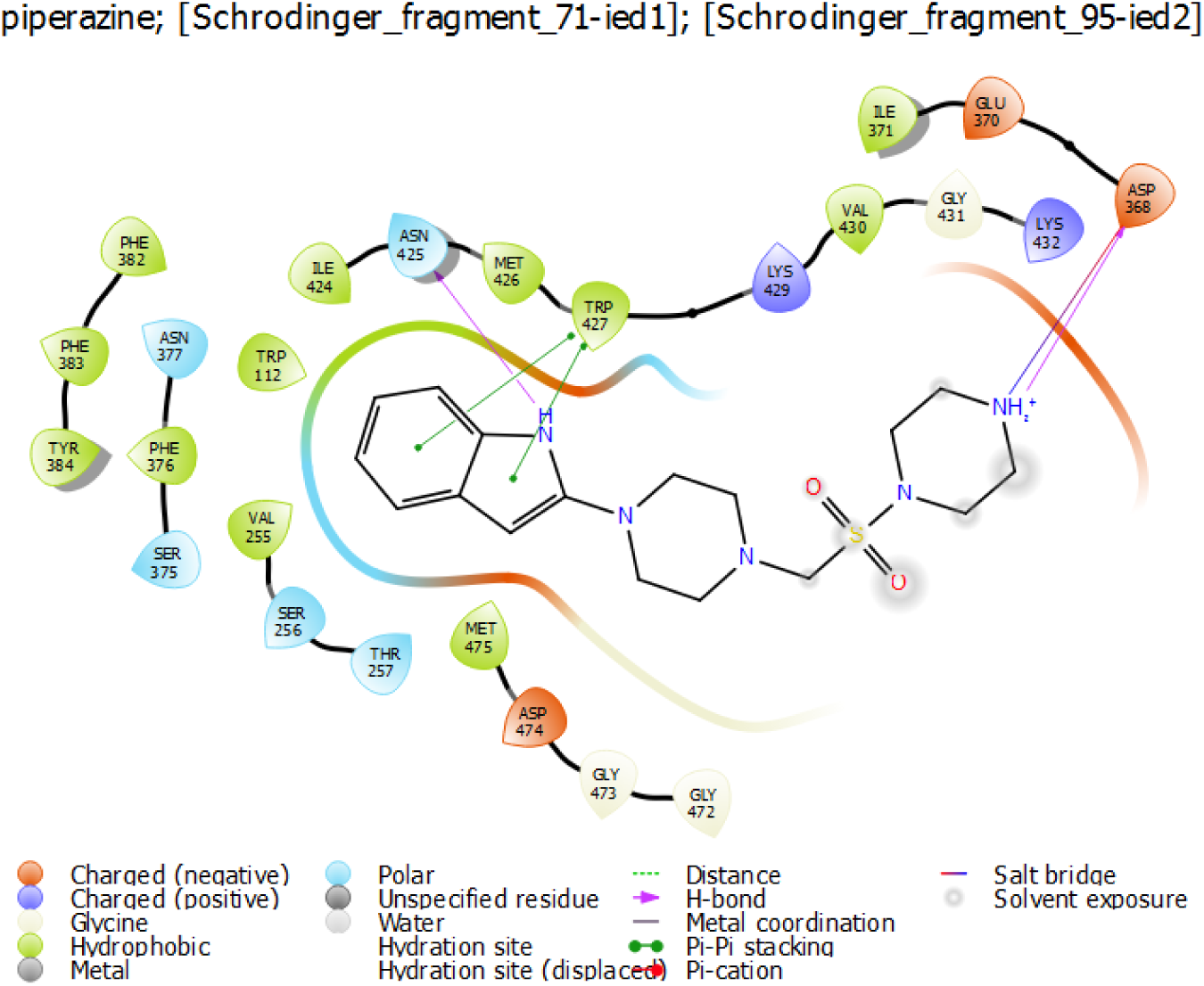
IFD interaction diagram for ligand 059.

**Figure 9:**
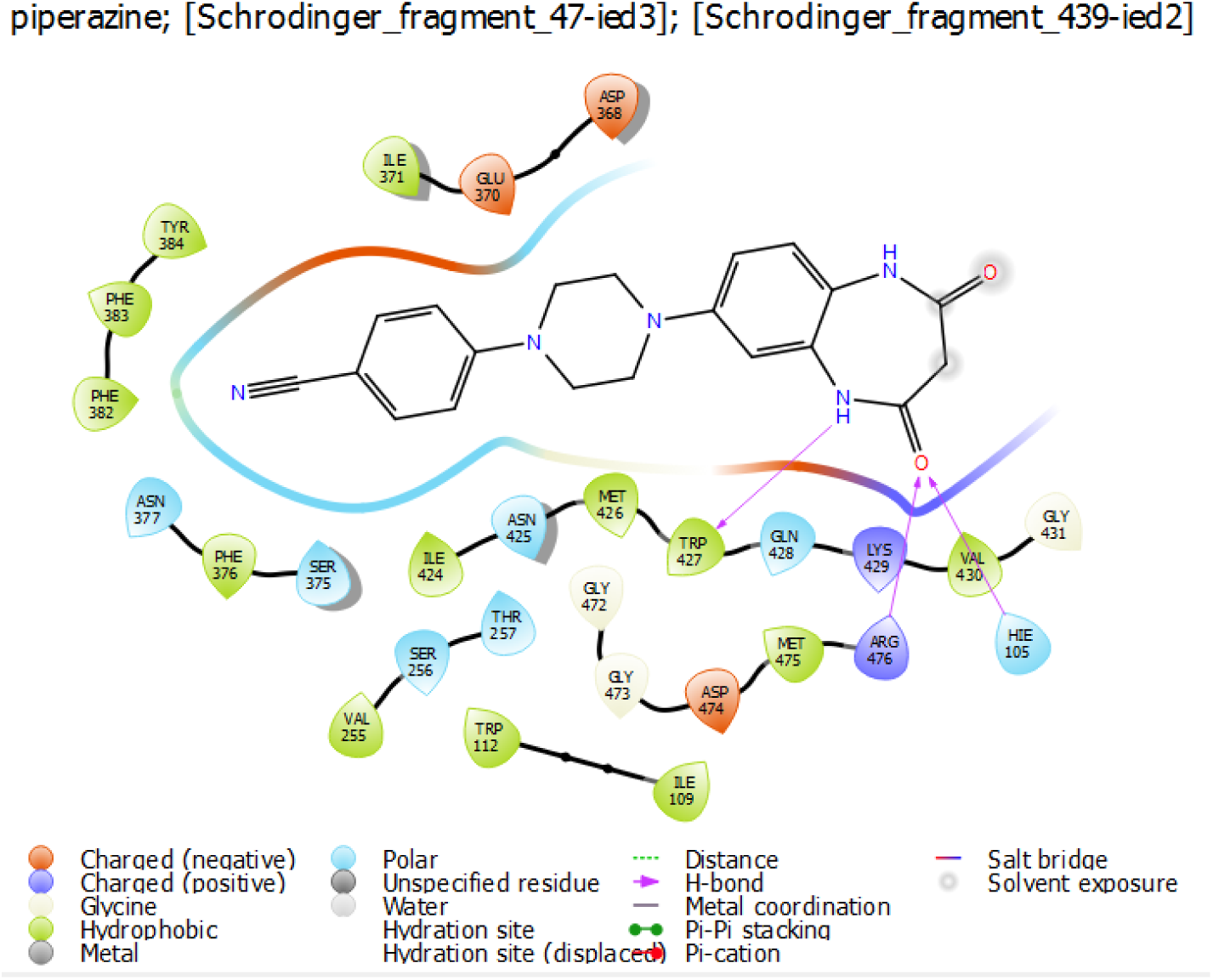
IFD interaction diagram for ligand 177.

**Figure 10:**
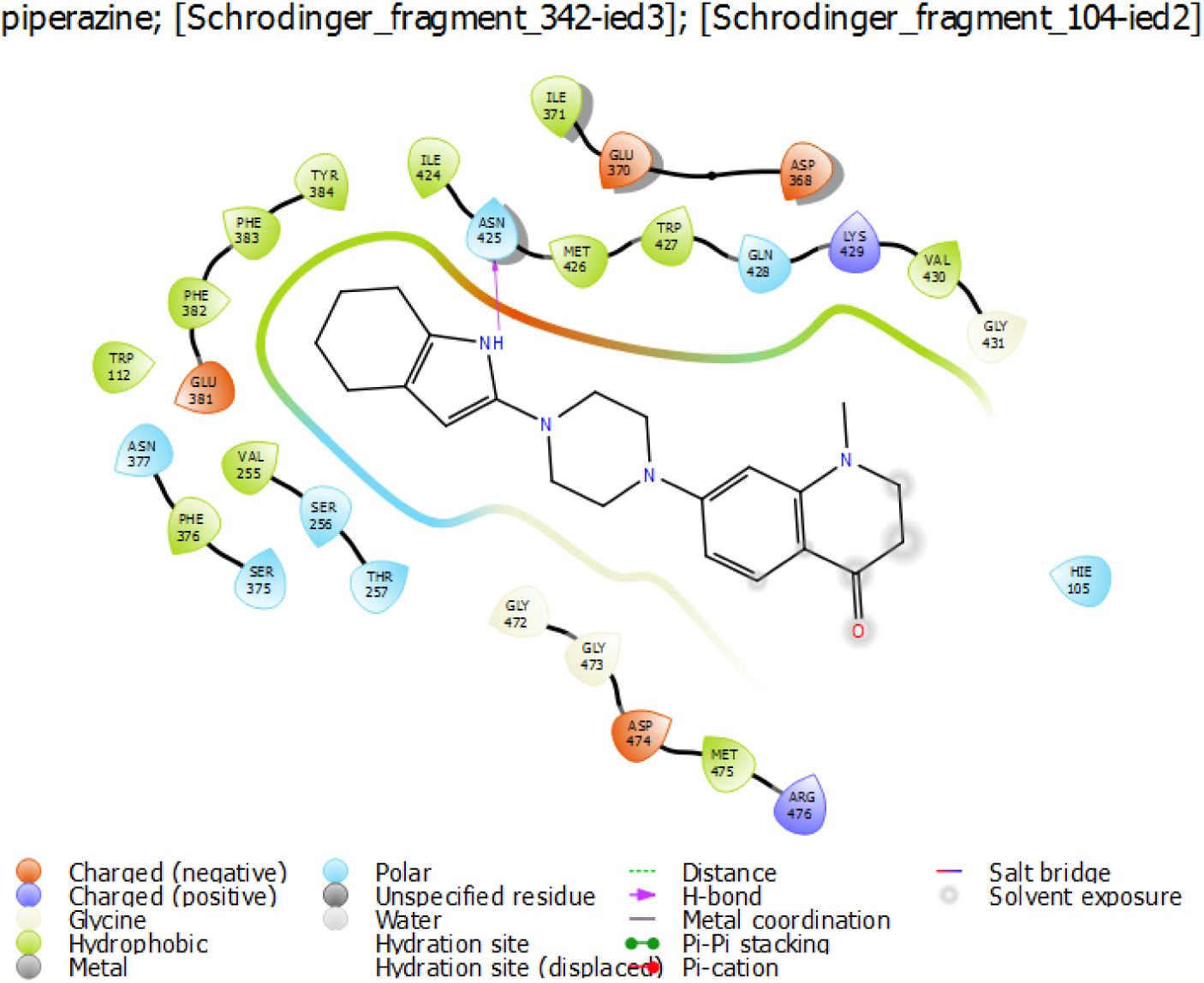
IFD interaction diagram for ligand 085.

**Figure 11:**
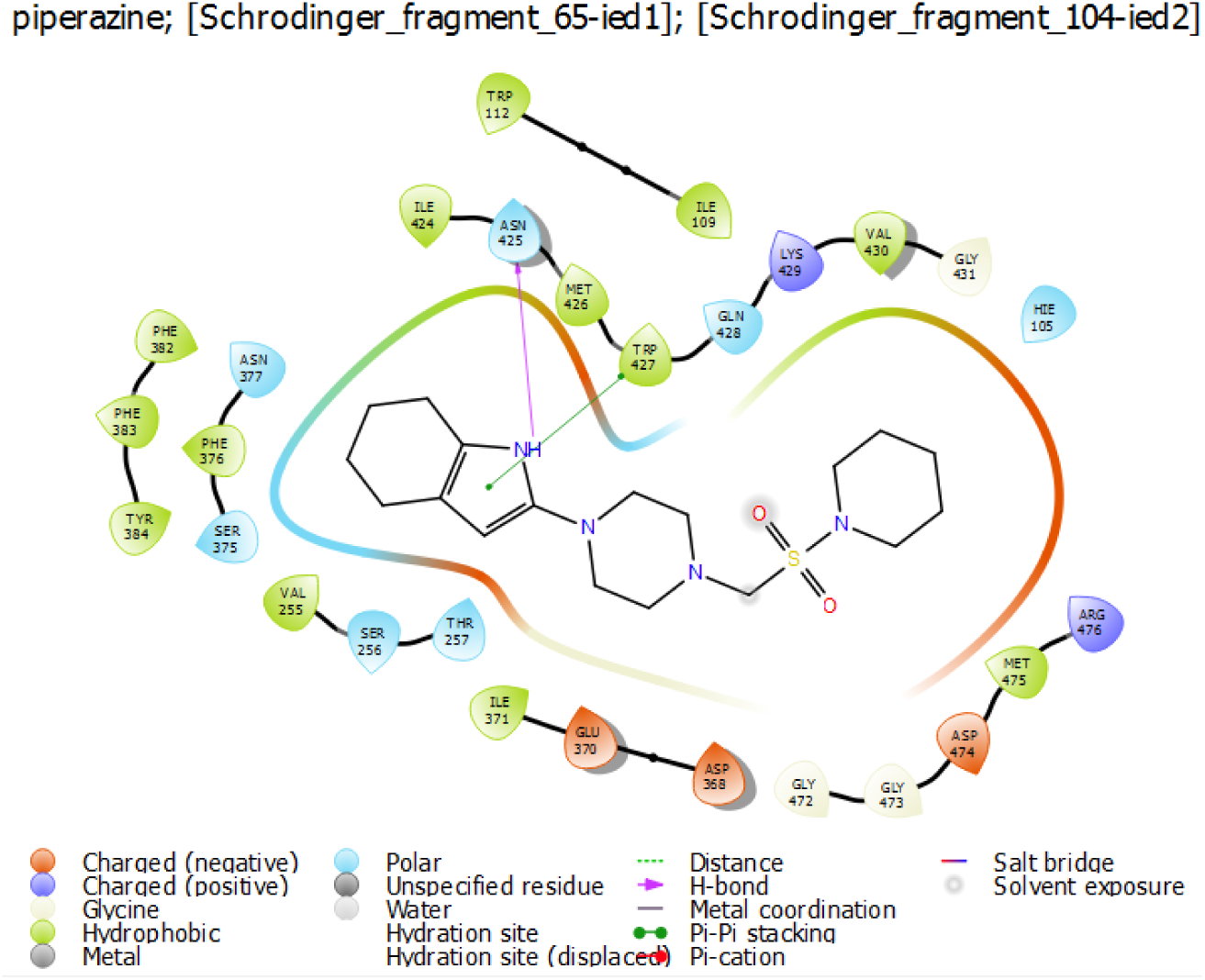
IFD interaction diagram for ligand 044.

**Figure 12:**
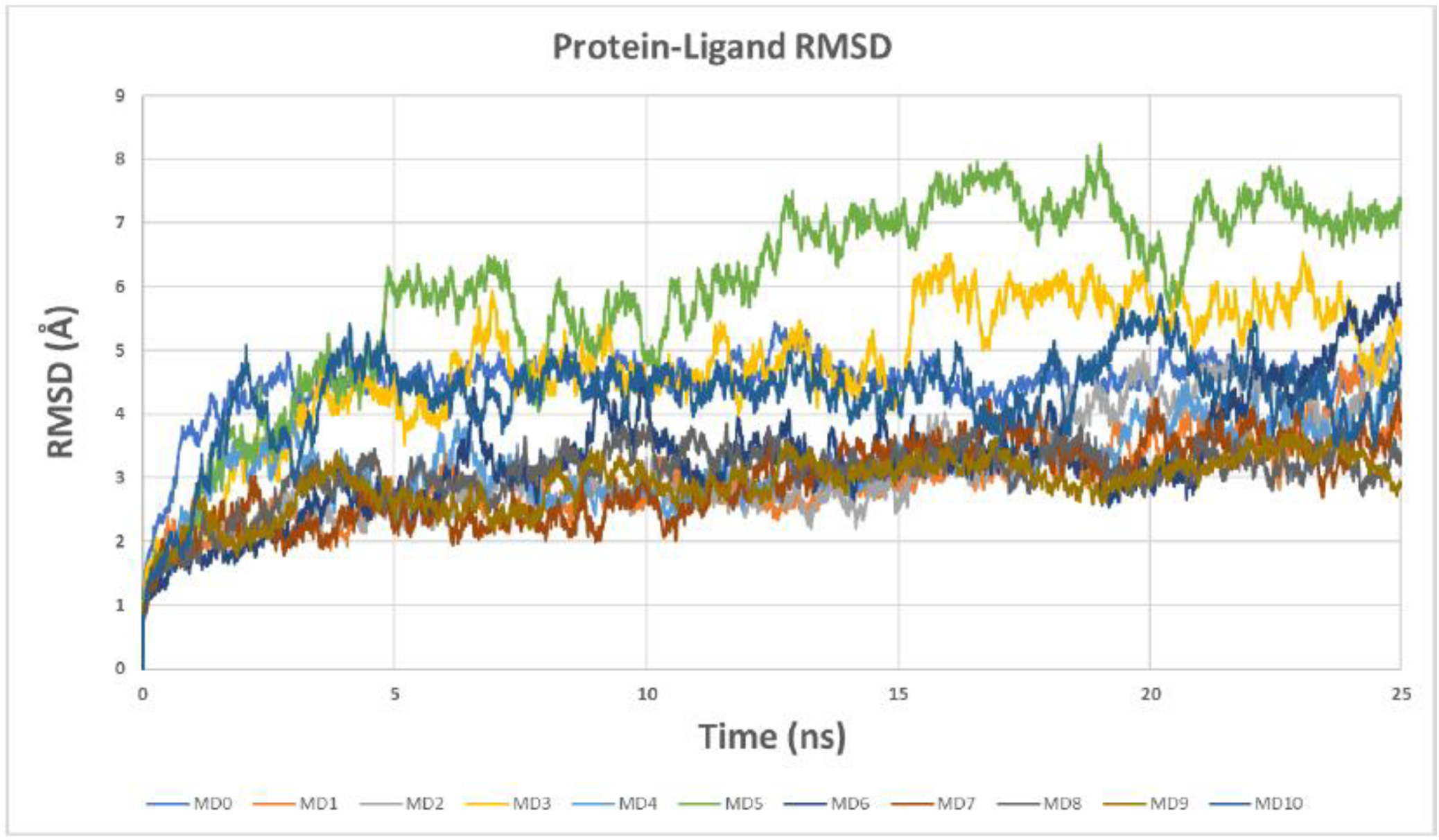
Protein-ligand root-mean-square deviation for all molecular dynamics simulations performed.

**Figure 13:**
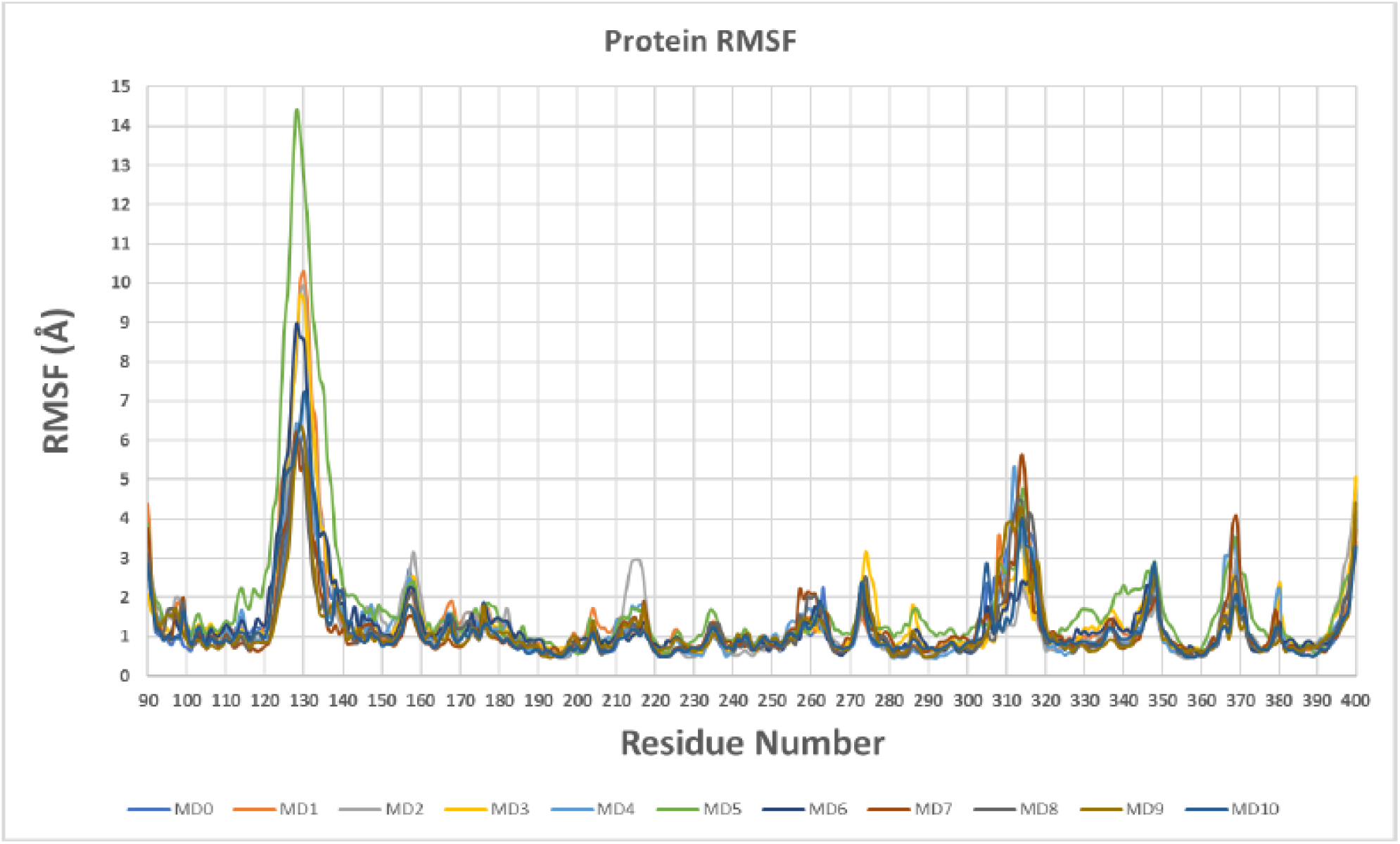
Protein-ligand root-mean-square fluctuation for all molecular dynamics simulations performed.

